# Establishment of an efficient one-step enzymatic synthesis of cyclic-2,3-diphosphoglycerate

**DOI:** 10.1101/2025.02.21.639460

**Authors:** Christina Stracke, Benjamin H. Meyer, Simone A. DeRose, Erica Elisa Ferrandi, Ilya V. Kublanov, Michail N. Isupov, Nicholas J. Harmer, Daniela Monti, Jennifer Littlechild, Felix Müller, Jacky L. Snoep, Christopher Bräsen, Bettina Siebers

## Abstract

Extremolytes – unique compatible solutes produced by extremophiles - protect biological structures like membranes, proteins, and DNA under extreme conditions, including extremes of temperature and osmotic stress. These compounds hold significant potential for applications in pharmaceuticals, healthcare, cosmetics, and life sciences. However, despite their promise, only a few extremolytes, such as ectoine and hydroxyectoine, are commercially established, primarily due to the lack of efficient production strategies for other compounds.

Cyclic 2,3-diphosphoglycerate (cDPG), a unique metabolite found in certain hyperthermophilic methanogenic Archaea, plays a key role in thermoprotection and is synthesized from 2-phosphoglycerate (2PG) through a two-step enzymatic process involving 2-phosphoglycerate kinase (2PGK) and cyclic-2,3-diphosphoglycerate synthetase (cDPGS). In this study, we present the development of an efficient *in vitro* enzymatic approach for the production of cDPG directly from 2,3-diphosphoglycerate (2,3DPG), leveraging the activity of the cDPGS from *Methanothermus fervidus* (*Mf*cDPGS).

We optimized the heterologous production of *Mf*cDPGS in *Escherichia coli* by refining codon usage and expression conditions. The purification process was significantly streamlined through an optimized heat precipitation step, coupled with effective stabilization of *Mf*cDPGS for both usage and storage by incorporating KCl, Mg^2+^, reducing agents and omission of an affinity tag. The recombinant *Mf*cDPGS showed a *V*_*max*_ of 38.2 U mg^-1^, with *K*_*M*_ values of 1.52 mM for 2,3DPG and 0.55 mM for ATP. The enzyme efficiently catalyzed the complete conversion of 2,3DPG to cDPG. Remarkably, even at a scale of 100 mM, it achieved full conversion of 37.6 mg of 2,3DPG to cDPG within 180 minutes, using just 0.5 U of recombinant *Mf*cDPGS at 55°C. These results highlight that *Mf*cDPGS can be easily produced, rapidly purified, and sufficiently stabilized while delivering excellent conversion efficiency for cDPG synthesis as value-added product. Additionally, a kinetic model for *Mf*cDPGS activity was developed, providing a crucial tool to simulate and scale up cDPG production for industrial applications. This streamlined process offers significant advantages for the scalable synthesis of cDPG, paving the way for further biochemical and industrial applications of this extremolyte.

## 1 Introduction

Compatible solutes are ubiquitous and distributed across all three domains of life, accumulating within cells to high concentrations (up to 2 M) in response to diverse environmental stressors such as heat, cold, osmotic stress, and desiccation. These solutes integrate seamlessly with cellular metabolism, enabling organisms to thrive under harsh conditions by stabilizing and protecting nucleic acids, proteins and cellular structures (da Costa, Santos, and Galinski 1998; Lamosa et al. 1998). Cells either acquire compatible solutes from their environment or produce them through *de-novo* synthesis. While their diversity is vast, compatible solutes generally fall into a few different chemical categories, including (i) amino acids (e.g., α-glutamate, β-glutamate) and amino acid derivatives (e.g., glycine-betaine, ectoine, hydroxyectoine) and (ii) sugars (e.g. trehalose, mannosylglycerate), polyols (e.g., glycerol) and their derivatives (Empadinhas and da Costa 2006). Their remarkable protective properties for enzymes, DNA, membranes and entire cells have led to their incorporation into various commercial applications across industries such as food, health and consumer care, and cosmetics (Becker and Wittmann 2020). Extremolytes - compatible solutes synthesized by extremophiles such as halophiles or (hyper)thermophiles - have garnered significant industrial interest. Beyond the ubiquitous, widely utilized compatible solute trehalose, the most commercially prominent extremolytes are ectoine and hydroxyectoine (Lentzen and Schwarz 2006b, Becker and Wittmann 2020). Additionally, phosphorylated compounds have been identified as extremolytes in hyperthermophiles, including di-*myo*-1,1’-inositol-phosphate (DIP) (Scholz et al. 1992; Martins and Santos 1995), α-diglycerol phosphate (DGP) (Borges et al. 2006), and cyclic 2,3-diphosphoglycerate (cDPG) (Seely and Fahrney 1983; Hensel and König 1988; Lehmacher, Vogt, and Hensel 1990). However, despite their promising properties, these phosphorylated extremolytes remain largely untapped for biotechnological and industrial applications, primarily due to the lack of suitable production hosts and cost-effective synthesis strategies. The extremolyte cDPG has been uniquely reported in the hyperthermophilic Methanoarchaea, such as *Methanothermus fervidus, Methanopyrus kandleri* and *Methanothermobacter thermoautotrophicus*, at concentrations ranging from 0.3 - 1.1 M (Hensel and König 1988; Ciulla et al. 1994; Matussek et al. 1998; Shima et al. 1998). It contributes to protein thermoprotection and has been shown to enhance the thermostability of various model enzymes (Hensel and Jakob 1993; Shima et al. 1998; Borges et al. 2002). Additionally, cDPG protects plasmid DNA from oxidative damage by hydroxyl radicals and acts as a superoxide scavenger (Lentzen and Schwarz 2006b).

The enzymatic synthesis of cDPG involves two distinct catalytic steps. First, 2-phosphoglycerate kinase (2PGK) catalyzes the conversion of 2-phosphoglycerate (2PG) into the phosphate diester 2,3-diphosphoglycerate (2,3DPG). In the subsequent step, cyclic-2,3-diphosphoglycerate synthetase (cDPGS) forms an intramolecular phosphoanhydride bond, yielding cDPG (Figure 1A). Despite the well-documented protective properties of cDPG and its industrial potential, an efficient production process remains elusive. This challenge stems from several factors, including the low cell yields and demanding cultivation requirements of natural producer strains, such as the obligate anaerobic growth conditions, high temperature and salt requirements of this methanogens. In addition, the lack of genetic tools for these organisms make a whole-cell *in vivo* approach for cDPG production in natural hosts impractical.

**Figure 1:**
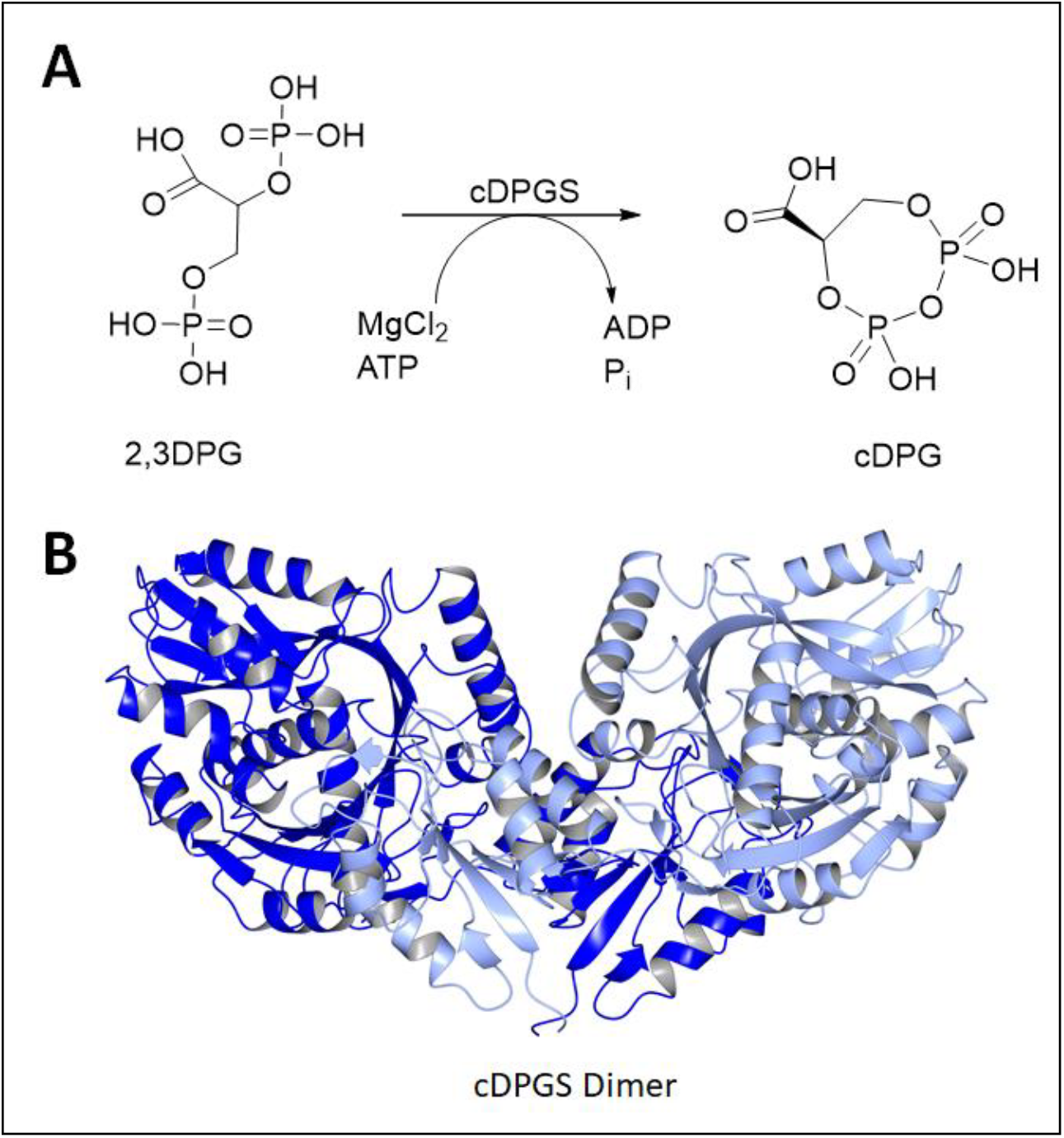
Pathway for cDPG synthesis and structure of *Mf*cDPGS dimer. **(A)** Schematic representation of the enzymatic synthesis of cyclic 2,3-diphosphoglycerate (cDPG) by the formation of an intramolecular phosphoanhydride bond in 2,3-diphosphoglycerate (2,3DPG). **(B)** Enzyme structure of the *Methanothermus fervidus* cDPGS dimer (PDB 8ORU viewed perpendicular to its molecular dyad. The two subunits are shown in blue and light blue (visualized using CCP4mg (McNicholas et al. 2011)).

In a previous study, we successfully employed an *in vivo* metabolic engineering approach to produce cDPG in the thermophilic bacterium *Thermus thermophilus*. By introducing the *Mf*2PGK and *Mf*cDPGS from *M. fervidus*, into *T. thermophilus* HB27, we achieved *in vivo* production of 650 µM cDPG within 16 hours (De Rose et al., 2021). More recently, we solved the crystal structure of the *Mf*cDPGS (PDB 8ORK, Figure 1B), including its complex with 2,3DPG and ADP (PDB 8ORU), offering novel mechanistic insights (De Rose et al., 2023). Notably, the *Mf*cDPGS features a unique N-terminal domain (127 amino acids) with a novel fold, which is essential for stabilizing the active dimeric structure and facilitating ligand interactions (De Rose et al., 2023). In this study as an alternative approach to *in vivo* production in *T. thermophilus*, we explored an *in vitro* enzymatic approach for large-scale synthesis of cDPG. Therefore, to improve the efficiency of heterologous *Mf*cDPGS expression in *Escherichia coli*, we applied codon optimization, omitted the affinity tag, refined expression conditions, and streamlined the purification process by incorporating a simple heat precipitation step. Additionally, we optimized the stabilization conditions of the recombinant protein for both immediate use and long-term storage. Finally, we demonstrated complete substrate conversion to the product, precisely described by our mathematical model, enabling straightforward upscaling and downstream processing of cDPG for future applications.

## 2 Material and Methods

### 2.1 Cloning, heterologous expression and purification of the recombinant proteins

#### Cloning

Initially the cyclic 2,3-diphosphoglycerate synthetase (*cdpgs*, 1383 bp) from *M. fervidus* (*Mfe*r_0077, CAA70986) was amplified via PCR from chromosomal DNA of *M. fervidus* (for primers see Supplementary Table 1). The PCR product was purified using the Wizard^®^ SV Gel and PCR Clean-Up kit (Promega, Fitchburg, USA) according to the instructions of the manufacturer. Restricted DNA fragments were cloned into the expression vector pET24a (Kan^R^) (Merck, Darmstadt, Germany) with C-terminal His-tag and were transformed into*E. coli* DH5α (Life Technologies, Carlsbad, USA). Later, the *cdpgs* gene was codon optimized (Eurofins, Ebersberg, Germany) for expression in *E. coli* and subcloned in pET15b (Amp^R^) (Merck, Darmstadt, Germany) either with N-terminal His-tag or without an affinity tag (for primers see Supplementary Table 1). All sequences and plasmids were verified by sequencing (LGC genomics, Berlin, Germany).

#### Heterologous expression

Competent cells of various *E. coli* strains (BL21(DE3) and BL21(DE3) CodonPlus-pRIL (Cam^R^) (Merck, Darmstadt, Germany), Rosetta (DE3) (Cam^R^) (Stratagene, San Diego, USA) were transformed with the constructed plasmids and grown in 200 mL – 400 mL lysogeny broth medium (LB broth) (Carl Roth GmbH + Co. KG, Karlsruhe, Germany) at different expression temperatures (37°C, 30°C, 16°C). The expression was induced by the addition of 1 mM isopropyl-β-d-thiogalactopyranoside (IPTG) at an OD_600_ between 0.6 - 0.8. Cultures were incubated in a Unitron shaker (Infors, Basel, Switzerland) over night at 180 rpm. Cells were harvested (6 000 x g, 20 min, 4°C) and 1 g of cells (wet weight) were resuspended in 5 mL of resuspension buffer (50 mM potassium phosphate (K_2_HPO_4_/KH_2_PO_4_) (pH 8), 2.5 mM MgCl_2_, 5 mM DTT, 300 mM KCl for affinity chromatography of His-tagged *Mf*cDPGS and 50 mM 2-(N-morpholino)ethanesulfonic acid (MES)/KOH (pH 6.5), 10 mM DTT, 10 mM MgCl_2_ , 400 mM KCl for *Mf*cDPGS without affinity-tag. For cell lysis, cells were passed three times through a French pressure cell at 20 000 psi. Cell debris was removed by centrifugation (25 000 x g, 45 min, 4°C).

#### Purification

The crude extract containing the recombinant thermophilic *Mf*cDPGS was subjected to a heat precipitation step at 75°C for 30 min in an Eppendorf Thermomixer at 700 rpm (Eppendorf, Hamburg, Germany) followed by centrifugation (25.000xg, 45 min, 4°C) to remove denatured mesophilic host proteins. The His-tagged *Mf*cDPGS was purified using affinity chromatography via Protino^®^ Ni-tris-carboxymethyl ethylene diamine (TED) columns (0.25 g columns for 9.6 mg (1.37 mg/ml) and 0.5 g columns for 20.8 mg (4.15 mg/mL) soluble fraction after heat precipitation) (Macherey-Nagel GmbH & Co. KG, Düren, Germany). The manufacturer’s instructions were followed with the exception of buffer composition during equilibration and washing. Instead of the supplied 50 mM sodium phosphate (NaH_2_PO_4_) (pH 8.0), 300 mM NaCl, 50 mM potassium phosphate (K_2_HPO_4_/KH_2_PO_4_) (pH 8), 2.5 mM MgCl_2_, 5 mM DTT, 300 mM KCl was used. This adjustment was made due to the methanogenic origin of the *Mf*cDPGS, as the intracellular K^+^ concentration of *M. fervidus* ranges from 500 to 900 mM under optimal growth conditions, much higher than that of mesophilic methanogens or other thermophilic archaea (Hensel & König, 1998). For elution, the potassium phosphate buffer was supplemented with 250 mM imidazole. Elution fractions containing *Mf*cDPGS activity were pooled and dialyzed using Spectra/Por^®^ standard RC dialysis membrane (Repligen, Waltham, USA) with a molecular weight cutoff of 30 kDa. For the recombinant *Mf*cDPGS without an affinity tag, the protein purification was further modified. The crude extract (13.1 mg/mL) was diluted with 50 mM MES/KOH (pH 6.5), 10 mM DTT, 10 mM MgCl_2_, and 400 mM KCl prior to the heat precipitation step. Dilutions of 1:2 (∼ 6.6 mg/ml), 1:4 (∼ 3.3 mg/mL), 1:6 (∼ 2.2 mg/mL) were tested and incubated at 75°C for 30 min in an Eppendorf Thermomixer at 700 rpm (Eppendorf, Hamburg, Germany), followed by centrifugation (25.000xg, 45 min, 4°C) to remove denatured mesophilic host proteins. For further experiments a dilution at a 1:6 ratio was used.

Additional purification was achieved by size exclusion chromatography (SEC). Therefore, the soluble fraction after heat precipitation was concentrated using a Vivaspin^®^ concentrator (30 kDa cutoff, Satorius, Göttingen, Germany), filtered (0.45 µm polyvinylidene fluoride membrane, Carl Roth, Germany), and 5 mg of protein were loaded onto a Superose Increase 10/300 column (Cytiva Life Sciences, Freiburg, Germany). The column (24 ml volume, flow rate 0.5 mL/min) was equilibrated in 50 mM MES/KOH (pH 6.5), 400 mM KCl. Protein expression, purification and subunit molecular mass of *Mf*cDPGS were analyzed by denaturing SDS-polyacrylamide gel electrophoresis. All protein concentrations were determined using a modified Bradford method (Zor and Selinger 1996) with bovine serum albumin as a standard (Bio-Rad, Feldkirchen, Germany).

### 2.2 *M. fervidus* cDPGS activity

#### Continuous PK-LDH assay

The enzymatic activity of *Mf*cDPGSwas determined at 55°C by coupling the ADP formation from ATP to the oxidation of NADH, using pyruvate kinase (PK) and _L_-lactate dehydrogenase (LDH) from rabbit muscle (Merck, Darmstadt, Germany) as auxiliary enzymes. The assay mixtures (total volume 0.5 mL) contained 50 mM MES/KOH (pH 6.5) with 10 mM MgCl_2_, 400 mM KCl, 2.5 mM ATP, 0.5 mM NADH, 2 mM phosphoenolpyruvate (PEP), 8 U PK, 4 U LDH and 4.8 µg purified *Mf*cDPGS (0.69 mg/mL). After 1 min preincubation at 55°C the reactions were started by the addition of 2,3DPG and the oxidation of NADH was followed in a Specord UV/VIS spectrometer (Analytic Jena AG, Jena, Germany) at 340 nm (extinction coefficient 6.22 mM^-1^ cm^-1^). To determine the kinetic constants for 2,3DPG and ATP, their concentrations were varied between 0 mM and 10 mM, while maintaining a Mg^2+^/ATP molar ratio of 0.5. Experimental data were fitted, and kinetic constants were determined using the NonlinearModelFit function in Wolfram Mathematica with a two substrate irreversible Michaelis Menten equation. The substrate 2,3DPG was used either as penta-sodium salt or as penta-cyclohexylammonium-salt (Merck, Darmstadt, Germany). It was ensured that the coupling enzymes were not rate limiting. One unit (1 U) of enzyme activity is defined as 1 µmol substrate of product (ADP) formed per minute. All measurements were performed in triplicates.

#### Effect of salt and pH

The effect of salt on *Mf*cDPGS enzyme activity was analyzed in the presence of KCl using the continuous PK-LDH assay at 55°C, with varying salt concentrations ranging from 0 to 400 mM KCl. The pH optimum was determined over a range from pH 5.5 to 8.0, using the following buffers: 50 mM MES/KOH (pH 5.5, 6.5), 50 mM potassium phosphate buffer (pH 6.8) and 50 mM HEPES/KOH (pH 7.0 and 8.0), all buffers were supplemented with 400 mM KCl.

#### Optimal Mg^2+^/ATP ratio

To determine the optimal molar ratio of Mg^2+^/ATP for the synthesis of cDPG, the formation of ADP from ATP was coupled to the oxidation of NADH via the continuous PK-LDH assay at 55°C using 50 mM MES/KOH (pH 6.5), 400 mM KCl, 10 mM DTT, 10 mM 2,3DPG and 4.8 µg of the purified *Mf*cDPGS (0.69 mg/mL). The assays were performed in presence of different molar ratios of Mg^2+^/ATP (a concentration range of 1 – 50 mM was investigated, corresponding to a molar ratio of 0.02 – 1.0). It was ensured that the auxiliary enzymes as well as 2,3DPG were not rate limiting. All experiments were performed in triplicates.

#### Long-term stability

The stability of purified *Mf*cDPGS was analyzed over a period of several month. To initiate the analysis, the enzyme stocks stored at -70°C was carefully thawed on ice and mixed thoroughly by pipetting. Subsequently, the residual enzymatic activity was measured using the continuous PK-LDH assay at 55°C.

#### Discontinuous assays for analysis of 2,3DPG conversion to cDPG

To determine the conversion of 2,3DPG (1 – 100 mM) to cDPG over time, discontinuous enzyme assays were performed at 55°C. The assays were conducted in a reaction buffer containing 50 mM MES/KOH (pH 6.5), 400 mM KCl, and 10 mM DTT. The molar ratio of 2,3DPG to ATP was maintained at 1:2, with 2,3DPG concentrations ranging from 1 to 100 mM and ATP concentrations from 2 to 200 mM. Additionally, the Mg^2+^/ATP ratio was adjusted between 0.2 and 0.5 to optimize enzyme activity. The reaction mixtures (1 mL) were incubated at 850 rpm and 55°C in an Eppendorf ThermoMixer^®^ (Eppendorf, Hamburg, Germany). Assays were performed in the presence of 0.5 U of the purified *Mf*cDPGS (0.49 mg/mL (1-10 mM conversions), 0.55 mg/mL (25 -100 mM conversions) from different enzyme purifications. At regular time intervals (0 min – 1250 min), 100 µL samples were withdrawn into pre-cooled reaction tubes and immediately stored at -70°C to halt the reaction. For the negative control, the same concentrations of 2,3DPG and ATP were used, but *Mf*cDPGS was omitted. ADP formation from ATP at each time point was quantified using the PK-LDH assay as an indicator reaction. The indicator reaction was performed at 25°C in a total volume of 200 µl in 96-well plates (Sarstedt, Nümbrecht, Germany) using a Tecan Microplate Reader at 340 nm (Tecan group Ltd., Lifesciences, Männedorf, Switzerland). To quantify the NADH formed, a standard curve (0 – 0.7 mM NADH) was used. To monitor the conversion of 2,3DPG to cDPG, aliquots from the enzyme assay were analyzed by ^31^P-NMR spectroscopy (Bruker Avance Neo 400 MHz, Bruker, Rheinstetten, Germany). Spectra were acquired in a H_2_O/D_2_O (8:2, v/v) mixture. Therefore, thawed samples from the respective time points were mixed with D_2_O (Merck, Darmstadt, Germany) and the assay buffer 50 mM MES/KOH (pH 6.5), 400 mM KCl). D_2_O was used in all experiments to provide a stable lock signal for the spectrometer (^1^H-NMR for NMR lock). The deuterium signal is used to stabilize the magnetic field, minimizing drift during long acquisitions and improving spectral quality. Spectra were processed, normalized and integrated using the TopSpin^©^ 3.6.2 software (Bruker BioSpin, Rheinstetten, Germany). As a positive control, for product validation 5 mM cDPG was solved in 50 mM MES/KOH (pH 6.5) supplemented with 400 mM KCl, 10 mM MgCl_2_, and 20% (v/v) D_2_O.

## 3 Results

### 3.1 Expression and purification of the *M. fervidus* cDPGS

Initially, the gene encoding *Mf*cDPGS (*cdpgs*, Mfer_0077, CAA70986) was cloned into pET24a vector with a C-terminal His-tag for recombinant expression in *E. coli* (Supplementary Figure S1). However, despite using various expression strains (*E. coli* BL21(DE3), BL21(DE3)-CodonPlus-pRIL (Merck, Darmstadt, Germany) and Rosetta (DE3) (Stratagene, San Diego, USA)), and lowering the expression temperature to 16°C, the expression of the tagged recombinant *Mf*cDPGS resulted only in low amounts of soluble protein. While small quantities were enriched in heat precipitation fractions (HP_75_ protein concentration = 1.37 mg/mL), the protein did not bind to the Ni-TED affinity column (Supplementary Figure S1). Hence, the *cdpgs* gene was further analyzed using the GenScript Rare Codon Analysis Tool (https://www.genscript.com/tools/rare-codon-analysis). The *cdpgs* sequence has a GC-content of 32.8%, and the Codon Adaption Index (CAI), which serves as a quantitative measure to predict gene expression levels in *E. coli*, yielded a value of only 0.54 (Supplementary Figure S2 & S3). Additionally, the Codon Frequency Distribution (CFD) analysis, which compares the distribution of codons in *cdpgs* with the typical codon usage in *E coli*, revealed 14 codons with frequencies below 31%, which are likely to impair expression efficiency in *E. coli* (Supplementary Figure S4). To address this, the *cdpgs* gene was synthesized as a codon optimized construct (Eurofins, Ebersberg, Germany, CAI = 0.75, GC-content 47.2%) for improved expression in *E. coli* (Supplementary Table S2). For recombinant expression, the codon-optimized *cdpgs* was subcloned in pET15b (with N-terminal His-tag) and expressed in *E. coli* BL21(DE3)-CodonPlus-pRIL. This approach yielded higher amounts of soluble protein (protein concentration after heat precipitation at 75°C (30 min) 4.15 mg/mL and after Ni-TED affinity chromatography 0.053 mg/mL) (Supplementary Figure S5). However, the obtained His-tagged *Mf*cDPGS was relatively unstable in solution and precipitated during dialysis to remove imidazole. To address this, we expressed the codon optimized gene without the affinity tag and reduced the expression temperature to 16°C. After overnight expression, the recombinant enzyme was purified to apparent homogeneity using a heat precipitation step (30 min at 75°C). We tested different protein concentrations (6.5 – 2.2 mg/mL), and 2.2 mg/mL yielded the best results for purity (Supplementary Figure S6). Size exclusion chromatography (SEC) was then used for further purification. From a 400 mL expression culture (1.7 g cell wet weight), 3.5 mg of recombinant *Mf*cDPGS was obtained after size exclusion chromatography. The molecular mass of the recombinant *Mf*cDPGS was determined to be 52 kDa under denaturing conditions by SDS-PAGE (Figure 2), which closely matches the calculated molecular mass of 50.7 kDa. The native molecular mass, determined by SEC was 119 kDa (Supplementary Figure S7), confirming the homodimeric structure of the protein, as observed in the crystal structure (De Rose et al., 2023).

**Figure 2:**
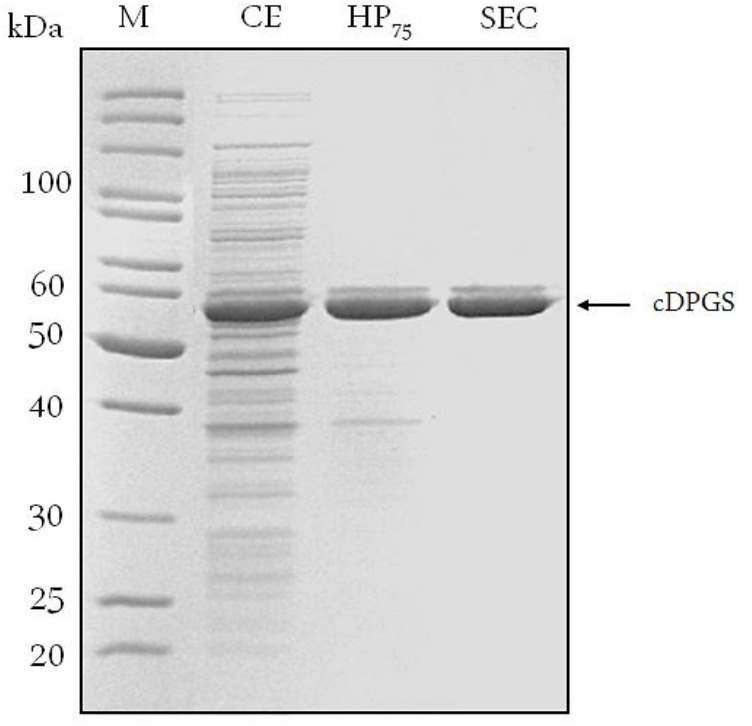
Purification of recombinant *Mf*cDPGS expressed in *E. coli*. The codon-optimized *cdpgs* gene was cloned into the pET15b vector without an affinity tag, and expression was optimized in *E. coli* BL21(DE3)-Codon-Plus. The recombinant protein was then purified by heat precipitation and size exclusion chromatography (SEC). Protein samples (5-10 µg) from each purification steps were analyzed by SDS-PAGE (12.5 %) and the gel was stained with Coomassie Brilliant Blue. CE: Crude extract, HP: Soluble fraction after heat precipitation (75°C, 30 min), SEC: Fraction after size exclusion chromatography. M: Protein marker, unstained protein ladder (Thermo Fisher Scientific, Carlsbad, USA).

### 3.2 Characterization of the *M. fervidus* cDPGS

*Mf*cDPGS catalyzes the ATP-depended formation of an intramolecular phosphoanhydride bond in 2,3DPG, resulting in cyclic 2,3-diphosphoglycerate (cDPG, Figure 1A). The enzyme activity of the purified *Mf*cDPGS (without a tag, after SEC) was determined at 55°C by following the ADP-formation from ATP using the continuous PK-LDH assay.

Preliminary tests (Mg^2+^/ATP ratio of 1.0) revealed that the activity of *Mf*cDPGS is highly salt-dependent, with a 5.5-fold activation observed in the presence of 400 mM KCl (Figure 3A). The optimal pH for *Mf*cDPGS activity was found to be at pH 6.5 (50 mM MES/KOH, 400 mM KCl) (Figure 3B). The optimal buffer conditions for enzyme characterization were identified as 50 mM MES/KOH (pH 6.5), supplemented with 400 mM KCl, 10 mM MgCl_2_ and 10 mM DTT. Notably, the inclusion of KCl, MgCl_2_ and DTT as reducing agent was essential throughout all purification steps. The rate dependence of the *Mf*cDPGS followed classical Michaelis-Menten kinetics, with a *V*_*max*_-value of 38.2 ± 1.7 U mg^-1^ and *K*_*M*_ -values of 1.52 ± 0.4 mM and 0.55 ± 0.08 mM for 2,3DPG and ATP, respectively (Figure 4). For long-term storage at -70°C, the addition of 25% (v/v) glycerol to the same buffer was necessary to preserve enzyme activity. After one week of storage at -70°C, the enzyme retained 100% of its activity. After 1.5 months, nearly full activity (95%) was maintained, and after 3.5-month 83% residual activity was observed (Figure 5).

**Figure 3:**
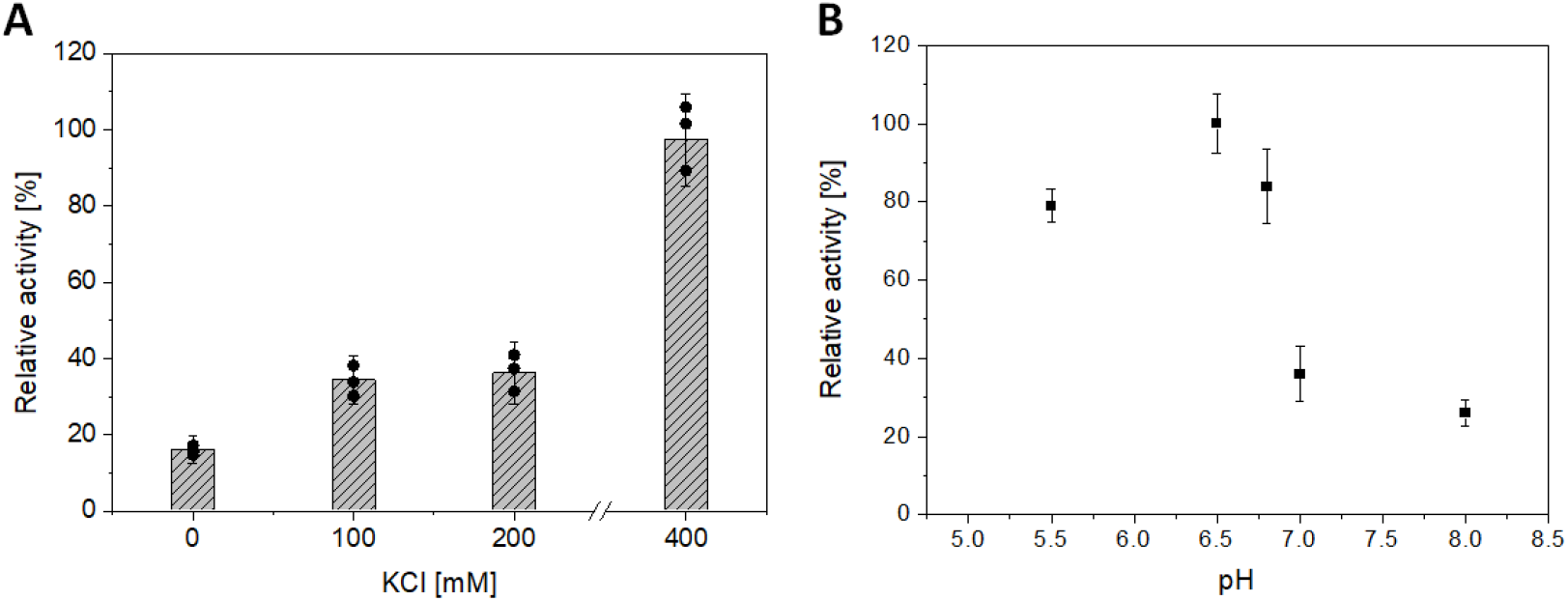
Effect of salt concentration and pH on the activity of *Mf*cDPGS. **(A)** Effect of KCl concentration (0 - 500 mM) and **(B)** effect of pH (5.5 - 8) on the activity of partially purified cDPGS (16.8 µg) after heat precipitation (75°C, 30 min). The enzyme activity was determined using the continuous PK-LDH assay at 55°C, which couples the formation of ADP from ATP to the oxidation of NADH monitored by the absorbance decrease at 340 nm. The pH optimum was determined using buffer systems as described in the Material and Method section, with each buffer supplemented with 400 mM KCl. The relative activity is expressed as a percentage, where 100 % corresponds to a specific activity of 1.8 U mg^-1^. All assays were performed in technical triplicates, and the error bars indicate the standard deviation of the mean.

**Figure 4:**
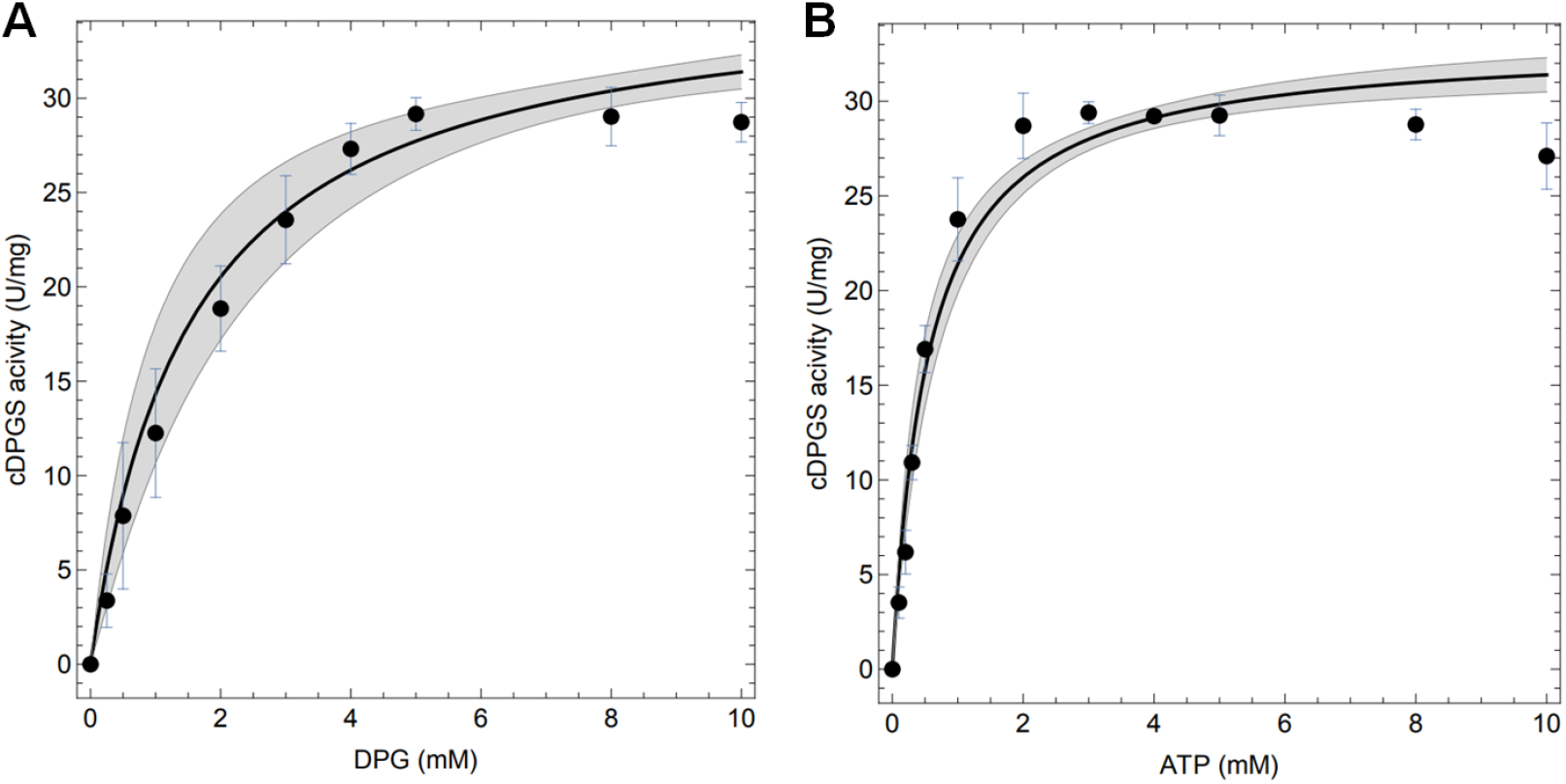
Enzymatic properties of the recombinant *Mf*cDPGS. Enzymatic activity **(A & B)** was measured at 55°C (340 nm) with 4.8 µg of purified enzyme using the continuous PK-LDH assay. **(A)** The specific activity of purified cDPGS was determined with varying concentration of 2,3 DPG (0-10 mM) and a constant 10 mM ATP. **(B)** The specific activity was determined with varying concentrations of ATP (0-10 mM) and a constant 10 mM 2,3-DPG. All assays were performed in technical triplicates, and the error bars represent the standard deviation of the mean. Computational fits to the complete dataset are shown as solid black lines and the mean prediction confidence intervals are represented as shaded areas.

**Figure 5:**
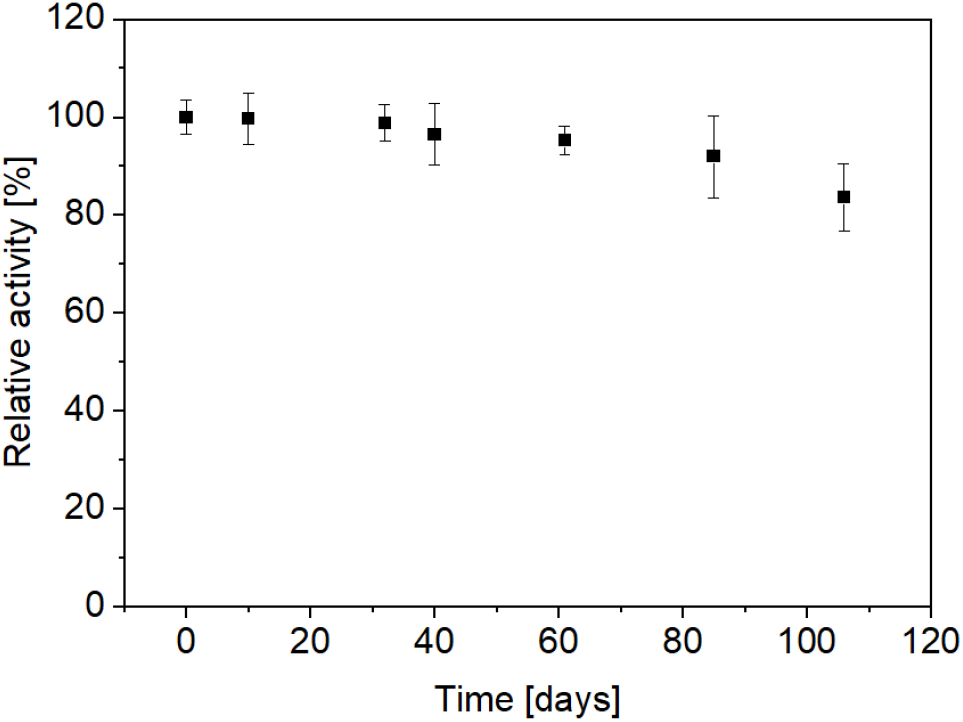
Long term stability of the recombinant *Mf*cDPGS. The enzyme was stored in aliquots at - 70°C in a buffer containing MES/KOH (pH 6.5), 25% (v/v) glycerol, 400 mM KCl, 10 mM MgCl_2_ and 10 mM DTT. Aliquots were thawed gently on ice and mixed thoroughly by pipetting before activity measurement. The residual activity of the thawed enzyme samples was determined at the indicated time points using the continuous PK-LDH assay. 100 % of relative activity corresponds to a specific activity of 18.7 U mg^-1^. All measurements were performed in technical triplicates, the error bars represent the standard deviation of the mean. Results are based on two biological replicates, each from a separate expression and purification run.

### 3.3 cDPG production

To determine the *in vitro* production of cDPG from 2,3DPG, we conducted discontinuous enzyme assays over a range of 2,3DPG concentrations (1-100 mM; 1:2 molar ratio of 2,3DPG to ATP) and analyzed the conversion efficiencies both enzymatically, using the discontinuous PK-LDH assay (for ADP determination), and *via* ^31^P-NMR spectroscopy (for 2,3DPG consumption and cDPG production). In initial experiments optimal Mg^2+^ /ATP ratios and product inhibition was investigated. To determine the optimal Mg^2+^/ATP ratio, a range of 0.02 – 1.0 (1 - 50 mM) was investigated, with the optimal ratio found to be between 0.2 - 0.5 (Supplementary Figure S8). Additionally, potential product inhibition by cDPG (0.5 – 20 mM) was assessed at a 1:2 molar ratio (5 mM 2,3DPG/10 mM ATP 10) and a 0.5 Mg^2+^/ATP ratio (5 mM Mg^2+^, 10 mM ATP) at 55°C. However, no significant inhibitory effects were observed (Supplementary Figure S9). Based on these findings, optimized *in vitro* conversions were performed using the following conditions: 50 mM MES/KOH (pH 6.5), 400 mM KCl, 10 mM DTT, a 1:2 molar ratio of 2,3 DPG/ATP, a Mg^2+^/ATP ratio of 0.2 -0.5, and 0.5 U of the purified recombinant *Mf*cDPGS. To monitor the conversion of 2,3DPG to cDPG, ^31^P-NMR analyses were conducted under these conditions. Samples from the discontinuous assays were mixed with 20% (v/v) D_2_O prior to analysis. A 5 mM cDPG standard was used for validation to facilitate accurate signal interpretation under these conditions (Supplementary Figure S10).

Finally, the conversion of 1 mM, 4 mM, 10 mM, 25 mM, 50 mM and 100 mM 2,3DPG to cDPG was monitored (1 mL reactions, 0.5 U *Mf*cDPGS) over time (0 -180 min, for 100 mM 2,3 DPG, also after 20 h) using both analytical approaches. Remarkably, complete conversion of 2,3DPG to cDPG was observed for all concentrations tested (Figure 6, and Supplementary Figure S11 A-F). ^31^P-NMR experiments showed full conversion of 1 mM, 4 mM, and 10 mM 2,3DPG after 25 - 40 min (Figure 6 A, and Supplementary Figure S12 A-C). For higher concentrations of 25 mM, 50 mM, and 100 mM 2,3 DPG, complete conversion was achieved after 180 min (Figure 6 B and Supplementary Figure S12 E-F). The enzymatic assay for ADP formation generally follows the same trend, although some deviations were observed due to experimental factors such as sample dilution.

**Figure 6:**
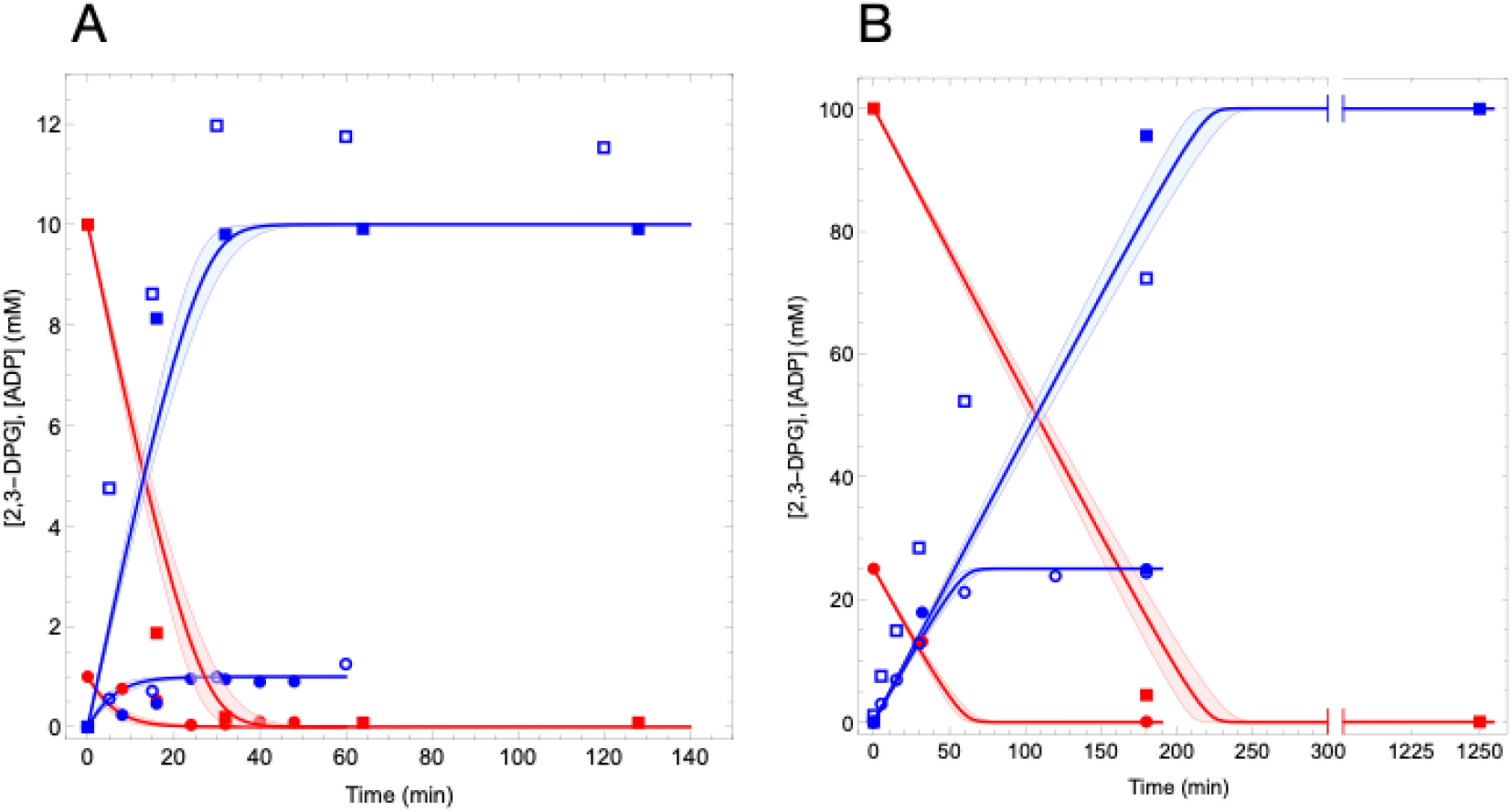
Conversion of 2,3 DPG to cDPG by recombinant *Mf*cDPGS. The time-dependent conversion results for (**A**) 1 mM (circles) and 10 mM (squares) conversion of 2,3 DPG and (**B**) 25 mM (circles) and 100 mM (squares) 2,3 DPG to cDPG are shown. Assays were performed at 55°C in 50 mM MES/KOH (pH 6.5), 400 mM KCl and a Mg^2+^/ATP ratio of 0.2 using 0.5 U of purified cDPGS. Samples were taken at regular time intervals, and resulting ADP from ATP was quantified using the PK-LDH assay and are shown with blue open symbols. Each data point represents the mean value of three independent technical replicates. The redfilled symbols represent the 2,3 DPG concentrations, while the blue filled symbols represent the cDPG concentrations, both quantified via ^31^P-NMR spectroscopy. Model (eq. 1 and 2) predictions for the experiments are shown with a red line for 2,3-DPG consumption and a blue line for cDPG production.

Furthermore, we simulated our experimental data with a computational model based on the kinetic parameters of *Mf*cDPGS. For the simulations we integrated the enzyme kinetic rate equation (eq. 1) as ordinary differential equations (eq. 2), using the experimentally determined kinetic parameters.

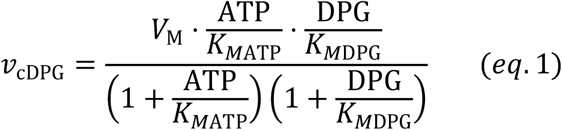

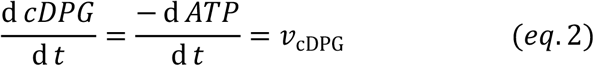

The model predicted the conversion rates for all experiments, aligning closely with our measured data (solid blue and red lines, Figure 6 A–B, and Supplementary Figure S11 A-F). The model also provided predictions for the time required to achieve complete conversion at various 2,3DPG concentrations: 1 mM (∼20 min), 4 mM (∼20 min), 10 mM (∼30 min), 25 mM (∼70 min), 50 mM (∼120 min), and 100 mM (∼200 min). Importantly, the model can be extended to predict conversion rates and outcomes for significantly larger production scales.

In conclusion, using the optimized protocol, the *Mf*cDPGS (0.5 U/mL) catalyzed the complete conversion of up to 100 mM 2,3DPG to cDPG in a total volume of 1 mL within approximately 180 min at 55°C, resulting in 37.6 mg of synthesized cDPG. Finally, to further streamline the enzymatic production of cDPG, we investigated whether the *Mf*cDPGS obtained after heat precipitation could be used directly for the conversion, thereby omitting the additional time- and cost-intensive purification step of SEC. The conversion of 10 mM 2,3DPG with 0.65 U/mL of heat-precipitated *Mf*cDPGS was monitored using ^31^P-NMR Complete substrate-to-product conversion was achieved within 16 min (Supplementary Figure S13). This simplified approach significantly reduces both protein production costs and process complexity, making it highly suitable for future applications.

## 4 Discussion

cDPG is a valuable extremolyte for diverse industrial and therapeutic applications. Despite its promising properties and potential applications, an efficient metabolic engineering strategy for cDPG production has not yet been established. A promising alternative to metabolic engineering is cell-free *in vitro* bioproduction using purified enzymes and cofactors in combination with enzyme kinetic modeling (Shen et al., 2020). This approach can circumvent challenges such as cellular regulation, competing metabolic pathways, kinetic bottlenecks, and toxicity issues (Fessner 2015, Claassens et al., 2019). However, efficient *in vitro* production depends on optimized enzyme expression and purification protocols, as well as strategies to ensure enzyme stability for prolonged use and storage.

To address these challenges, we optimized the recombinant production of the *M. fervidus* cDPGS and established a streamlined, one-step cell-free *in-vitro* production system for converting 2,3DPG to cDPG. Although the expression of the *Mf*cDPGS has been previously reported, earlier attempts at heterologous production yielded low expression levels and protein yields, with only 0.09 mg g^-1^ (wet weight) cells (Matussek et al. 1998). This limitation was also evident in our initial experiments.

Heterologous expression of methanogenic genes in non-methanogens has historically been challenging, often due to differences in gene structure and codon usage (Lange and Ahring 2001). Methanogens frequently use rare codons, such as AUA, AGA, and AGG, which are uncommon in

*E. coli*, leading to poor expression and low protein yields (Reeve 1993; Rosano and Ceccarelli 2009). Additionally, the prevalence of adenines (A) and thymines (T) at the third codon position in methanogenic genes further complicates expression in other host systems (Lange and Ahring 2001). Improved expression of archaeal methanogenic proteins in *E. coli* has been achieved in several instances through co-expression of rare tRNAs, such as argU, ileY, ileX, and leuW (Kim et al. 1998; Rosano and Ceccarelli 2009). To overcome these challenges, we opted to codon-optimize the *Mf*cDPGS-encoding gene for expression in *E. coli* and lowered the expression temperature, a widely used strategy for improving protein solubility and yield (Vera et al. 2007).

In addition to the expression challenges, the recombinant *Mf*cDPGS exhibited significant instability, consistent with previous studies on *Mf*cDPGS (Matussek et al. 1998; van Alebeek et al. 1994) and other methanogen-derived proteins (Breitung et al. 1992; Shima et al. 1998; Hensel and Jakob 1993; Liu et al. 2012; Reed et al. 2013). Furthermore, *Mf*cDPGS requires high concentrations of potassium ions (up 1 M) and magnesium ions (10 mM) for full activity (van Alebeek et al. 1994; Lehmacher and Hensel 1994; Lehmacher, Vogt, and Hensel 1990). Potassium ions and reducing agents like DTT also enhance enzyme stability and minimize oxygen susceptibility, as observed for other methanogens such as *M. kandleri* and *M. thermoautotrophicus* (Breitung et al. 1992; Liu et al. 2012; Reed et al. 2013). The primary challenge in this study was to optimize conditions for the expression, purification and stabilization of *Mf*cDPGS. We demonstrated that the enzyme’s activity was significantly enhanced by the addition of 400 mM KCl, 10 mM DTT, and 10 mM Mg^2+^. Furthermore, supplementing the storage buffer with 25% (v/v) glycerol enabled long-term storage at -70°C for several months, with minimal loss of activity.

Interestingly, the N-terminal His-tag introduced for purification appeared to exacerbate recombinant *Mf*cDPGS instability. This aligns with reports showing that affinity tags can sometimes interfere with protein function, folding, activity, and stability (Freydank, Brandt, and Dräger 2008; Young, Britton, and Robinson 2012; Booth et al. 2018). A different construct was used for optimal expression of the *Mf*cDPGS to obtain the optimal solubility and oligomeric homogeneity required for crystallization studies (De Rose et al., 2023).

The optimized procedure for *Mf*cDPGS production involved expressing the recombinant enzyme without any tags and performing all purification steps in the presence of 400 mM KCl, 10 mM MgCl_2_ and 10 mM DTT. The enzyme was efficiently purified using an optimized heat precipitation step. For further refinement, SEC could be added if required. This streamlined approach yielded a more than twenty-fold increase in protein recovery after SEC (2 mg g^-1^ cell wet weight) compared to previous reports, which achieved only 0.09 mg g^-1^ cells wet weight after affinity chromatography on reactive green 19 agarose (Matussek et al. 1998). A significant advantage of this method was maintaining a consistently high KCl concentration throughout the purification process. In contrast, the earlier method required low salt conditions during the affinity chromatography step, likely leading to substantial protein losses.

The recombinant *Mf*cDPGS produced by this new method exhibited markedly improved catalytic properties, with a significantly higher V_max_ of 38.2 U mg^-1^ at 55°C - more than double the previously reported value of 17 U mg^-1^ at 83°C (not considering the difference in assay temperature) - and closely approaching the activity described for the native enzyme (32 U mg^- 1^, 83°C) (Matussek et al. 1998). Additionally, the *K*_*M*_ -values for 2,3 DPG (1.52 mM) and ATP (0.55 mM) were significantly lower than those reported by Matussek and coworkers (*K*_*M*_ 2,3 DPG = 5.9 mM, *K*_*M*_ ATP= 3.0 mM) (Supplementary Table S3), likely reflecting differences in purification protocols and assay conditions between both studies.

The most significant outcome of this study was the demonstration that the recombinant *Mf*cDPGS catalyzed the complete conversion of 2,3 DPG to cDPG. In a 1 mL reaction volume, 100 µmol of substrate was fully converted within 180 min using 0.5 U of the purified enzyme, as confirmed by experimental data and computational modeling. The enzyme’s preference for the synthetic direction of the reaction is supported by thermodynamic considerations. The formation of the phosphoanhydride bond in 2,3DPG, driven by ATP hydrolysis, was shown to be thermodynamically favorable. While ATP hydrolysis to ADP and P_i_ has a ΔG^0’^ of -26 kJ mol^-1^, the hydrolysis of the anhydride bond in trianionic pyrophosphate and cDPG (Lentzen and Schwarz 2006b) has a calculated ΔG^0’^ of approximately -16 kJ mol^-1^ under standard conditions, (pH 7.0, ionic strength 0.1 M (eQuilibrator) (Flamholz et al. 2012). This suggests that the overall reaction is exergonic by approximately -10 kJ mol^-1^, providing a thermodynamic basis for the observed complete conversion.

The recently solved crystal structure of *Mf*cDPGS provides valuable insights into its substrate specificity and catalytic mechanism, further supporting the exergonic nature of the cyclisation reaction (De Rose et al. 2023). The structure reveals that *Mf*cDPGS remains in an “open” conformation until ATP binds correctly, triggering domain closure and creating a hydrophobic environment essential for the cyclisation reaction. Upon binding of ATP and 2,3 DPG, conformational changes position key residues in proximity, facilitating the catalytic process. the domain closure aligns the ATP γ-phosphate for a nucleophilic substitution (S_N_2) reaction, transferring the phosphate group to the 2-phophate of 2,3DPG. A second S_N_2 reaction then occurs, where the 3-phosphate of the intermediate attacks the 2-pyrophosphate, leading to the release of inorganic phosphate. This release of phosphate renders the reaction highly exergonic under standard conditions, providing the energy required for the formation of the phosphoanhydride bond in cDPG. (De Rose et al. 2023). These mechanistic insights underscore the enzyme’s efficiency and the thermodynamic feasibility of the reaction.

### 4.1 Conclusion

In conclusion, we have successfully developed an efficient protocol for the production and purification of *Mf*cDPGS to be used for cDPG production including strategies to stabilize the enzyme. Furthermore, we demonstrated the enzyme’s capability to achieve complete substrate conversion, enabling a streamlined and straightforward one-step biocatalytic synthesis of cDPG. This simplified approach enhances production efficiency and reduces downstream processing complexity. The established one-step *in vitro* synthesis is scalable and adaptable, with the potential to be extended into an enzyme cascade for the synthesis of cDPG from more cost-effective substrates, such as glycerate. These advancements pave the way for the cost-effective production of cDPG as a high-value product for diverse biotechnological applications.

## Abbreviations

cDPGS: cyclic-2,3-diphosphoglycerate synthetase
2,3DPG: 2,3-diphosphoglycerate
cDPG: cyclic 2,3-diphosphoglycerate
2PG: 2-phosphoglycerate
2PGK: 2-phosphoglycerate kinase

## 7 Conflict of Interest

The authors declare no competing interests.

### 8 Author Contributions

CS: Writing – original draft, Methodology, Investigation, Formal analysis, Validation. BHM: Writing – review & editing. SAR: Writing – review & editing. EEF: Writing – review & editing. IVK: Writing – review & editing. MNI: Writing – review & editing. NJH: Writing – review & editing. DM: Writing – review & editing. JAL: Writing – review & editing. FM: Writing – review & editing, Supervision. JLS: Writing – original draft, Software, Methodology. CB: Writing – original draft, Conceptualization, Methodology, Investigation, Formal analysis, Validation, BS: Writing – original draft, Methodology, Conceptualization, Project administration, Funding acquisition, Supervision, Data curation.

## 9 Acknowledgments

CS acknowledges funding by an Evonik Industries AG Scholarship. CS, BHM, CB and BS acknowledge funding by the German Federal Ministry of Education and Research (BMBF) grant HotSolute, 031B0612A within the ERA CoBioTech funding initiative. HotSolute has received funding from the European Union’s Horizon 2020 research and innovations programme under grant agreement No [722361]. JL, SAR, MNI, NJH acknowledge BBSRC BB/R02166X/1 grant for funding. JLS acknowledges funding from the DSI/NRF in South Africa (grant NRF-SARCHI-82813). DM acknowledges funding from the Italian Ministry of Education and Research (MIUR), Fondo per le agevolazioni alla ricerca “First 2016” (Decreto n. 110/2019). We thank Michaela Boraja for technical assistance. In addition, we would like to extend our gratitude to Bitop AG (Dortmund, Germany) for generously providing cDPG.

## References

Autengruber, A., U. Sydlik, M. Kroker, T. Hornstein, N. Ale-Agha, D. Stöckmann, A. Bilstein, C. Albrecht, A. Paunel-Görgülü, C. Suschek, J. Krutmann and K. Unfried (2014). “Signalling-Dependent Adverse Health Effects of Carbon Nanoparticles Are Prevented by the Compatible Solute Mannosylglycerate (Firoin) In Vitro and In Vivo.” PloS one 9: e111485.

Becker, J. and C. Wittmann (2020). “Microbial production of extremolytes — high-value active ingredients for nutrition, health care, and well-being.” Current Opinion in Biotechnology 65: 118–128.

Booth, W., C. Schlachter, S. Pote, N. Ussin, N. Mank, V. Klapper, L. Offermann, C. Tang, B. Hurlburt and M. Chruszcz (2018). “Impact of an N-terminal Polyhistidine Tag on Protein Thermal Stability.” ACS Omega 3: 760–768.

Borges, N., L. Gonçalves, M. Rodrigues, F. Siopa, R. Ventura, C. Maycock, P. Lamosa and H. Santos (2006). “Biosynthetic Pathways of Inositol and Glycerol Phosphodiesters Used by the Hyperthermophile Archaeoglobus fulgidus in Stress Adaptation.” Journal of bacteriology 188: 8128–8135.

Borges, N., C. D. Jorge, L. G. Gonçalves, S. Gonçalves, P. M. Matias and H. Santos (2014). “Mannosylglycerate: structural analysis of biosynthesis and evolutionary history.” Extremophiles 18(5): 835–852.

Borges, N., A. Ramos, N. D. Raven, R. J. Sharp and H. Santos (2002). “Comparative study of the thermostabilizing properties of mannosylglycerate and other compatible solutes on model enzymes.” Extremophiles 6(3): 209–216.

Breitung, J., G. Börner, S. Scholz, D. Linder, K. O. Stetter and R. K. Thauer (1992). “Salt dependence, kinetic properties and catalytic mechanism of N-formylmethanofuran:tetrahydromethanopterin formyltransferase from the extreme thermophile Methanopyrus kandleri.” Eur J Biochem 210(3): 971–981.

Cava, F., A. Hidalgo and J. Berenguer (2009). “Thermus thermophilus as biological model.” Extremophiles 13(2): 213–231.

Ciulla, R., C. Clougherty, N. Belay, S. Krishnan, C. Zhou, D. Byrd and M. F. Roberts (1994). “Halotolerance of Methanobacterium thermoautotrophicum delta H and Marburg.” J Bacteriol 176(11): 3177–3187.

Claassens, N. J., S. Burgener, B. Vögeli, T. J. Erb and A. Bar-Even (2019). “A critical comparison of cellular and cell-free bioproduction systems.” Curr Opin Biotechnol 60: 221–229.

Costa, J., N. Empadinhas, L. Gonçalves, P. Lamosa, H. Santos and M. S. da Costa (2006). “Characterization of the biosynthetic pathway of glucosylglycerate in the archaeon Methanococcoides burtonii.” J Bacteriol 188(3): 1022–1030.

da Costa, M., H. Santos and E. Galinski (1998). “An overview of the role and diversity of compatible solutes in Bacteria and Archaea.” Advances in biochemical engineering/biotechnology 61: 117–153.

De Rose, S., W. Finnigan, N. Harmer, J. Littlechild, T. consortium, S. Bettina, C. Bräsen, C. Stracke, M. Benjamin, I. N, H. J, A. Simone, A. Jennifer, B.-O. Elizaveta, S. Gavrilov, K. Ilya, D. Monti, E. Ferrandi, D. Eleonora and J. Snoep (2021). “Production of the Extremolyte Cyclic 2,3-Diphosphoglycerate Using Thermus thermophilus as a Whole-Cell Factory.” Frontiers in Catalysis 1: 803416.

De Rose, S. A., M. N. Isupov, H. L. Worthy, C. Stracke, N. J. Harmer, B. Siebers and J. A. Littlechild (2023). “Structural characterization of a novel cyclic 2,3-diphosphoglycerate synthetase involved in extremolyte production in the archaeon Methanothermus fervidus.” Front Microbiol 14: 1267570.

Empadinhas, N. and M. S. da Costa (2006). “Diversity and biosynthesis of compatible solutes in hyper/thermophiles.” Int Microbiol 9(3): 199–206.

Esteves, A. M., S. K. Chandrayan, P. M. McTernan, N. Borges, M. W. Adams and H. Santos (2014). “Mannosylglycerate and di-myo-inositol phosphate have interchangeable roles during adaptation of Pyrococcus furiosus to heat stress.” Appl Environ Microbiol 80(14): 4226–4233.

Faria, C., C. D. Jorge, N. Borges, S. Tenreiro, T. F. Outeiro and H. Santos (2013). “Inhibition of formation of α-synuclein inclusions by mannosylglycerate in a yeast model of Parkinson’s disease.” Biochim Biophys Acta 1830(8): 4065–4072.

Fessner, W. D. (2015). “Systems Biocatalysis: Development and engineering of cell-free “artificial metabolisms” for preparative multi-enzymatic synthesis.” N Biotechnol 32(6): 658–664.

Flamholz, A., E. Noor, A. Bar-Even and R. Milo (2012). “eQuilibrator-the biochemical thermodynamics calculator.” Nucleic Acids Res 40(Database issue): D770–775.

Freydank, A.-C., W. Brandt and B. Dräger (2008). “Protein structure modeling indicates hexahistidine-tag interference with enzyme activity.” Proteins: Structure, Function, and Bioinformatics 72(1): 173–183.

Göller, K. and E. A. Galinski (1999). “Protection of a model enzyme (lactate dehydrogenase) against heat, urea and freeze-thaw treatment by compatible solute additives.” Journal of Molecular Catalysis B: Enzymatic 7(1): 37–45.

Harishchandra, R. K., S. Wulff, G. Lentzen, T. Neuhaus and H. J. Galla (2010). “The effect of compatible solute ectoines on the structural organization of lipid monolayer and bilayer membranes.” Biophys Chem 150(1-3): 37–46.

Hensel, R. and I. Jakob (1993). “Stability of Glyceraldehyde-3-Phosphate Dehydrogenases from Hyperthermophilic Archaea at High Temperature.” Systematic and Applied Microbiology 16(4): 742–745.

Hensel, R. and H. König (1988). “Thermoadaptation of methanogenic bacteria by intracellular ion concentration.” FEMS Microbiology Letters 49(1): 75–79.

Huber, R., M. Kurr, H. W. Jannasch and K. O. Stetter (1989). “A novel group of abyssal methanogenic archaebacteria (Methanopyrus) growing at 110 °C.” Nature 342(6251): 833–834.

Kauth, M. and O. V. Trusova (2022). “Topical Ectoine Application in Children and Adults to Treat Inflammatory Diseases Associated with an Impaired Skin Barrier: A Systematic Review.” Dermatol Ther (Heidelb) 12(2): 295–313.

Kim, R., S. J. Sandler, S. Goldman, H. Yokota, A. J. Clark and S.-H. Kim (1998). “Overexpression of archaeal proteins in Escherichia coli.” Biotechnology Letters 20(3): 207–210.

Klähn, S., C. Steglich, W. R. Hess and M. Hagemann (2010). “Glucosylglycerate: a secondary compatible solute common to marine cyanobacteria from nitrogen-poor environments.” Environ Microbiol 12(1): 83–94.

Knapp, S., R. Ladenstein and E. A. Galinski (1999). “Extrinsic protein stabilization by the naturally occurring osmolytes beta-hydroxyectoine and betaine.” Extremophiles 3(3): 191–198.

Kumar, R., D. D. Patel, D. Bansal, S. Mishra, A. Mohammed, R. Arora Fls Frsc, A. Sharma, R. Sharma and R. Tripathi (2009). Extremophiles: Sustainable Resource of Natural Compounds-Extremolytes. Sustainable Biotechnology (pp. 279–294): 279-294.

Kunte, H., G. Lentzen and E. Galinski (2014). “Industrial Production of the Cell Protectant Ectoine: Protection Mechanisms, Processes, and Products.” Current Biotechnology 3.

Lamosa, P., L. Martins, M. da Costa and H. Santos (1998). “Effects of Temperature, Salinity, and Medium Composition on Compatible Solute Accumulation by Thermococcus spp.” Applied and Environmental Microbiology 64.

Lange, M. and B. K. Ahring (2001). “A comprehensive study into the molecular methodology and molecular biology of methanogenic Archaea.” FEMS Microbiol Rev 25(5): 553–571.

Lehmacher, A. and R. Hensel (1994). “Cloning, sequencing and expression of the gene encoding 2-phosphoglycerate kinase from Methanothermus fervidus.” Mol Gen Genet 242(2): 163–168.

Lehmacher, A., A. B. Vogt and R. Hensel (1990). “Biosynthesis of cyclic 2,3-diphosphoglycerate. Isolation and characterization of 2-phosphoglycerate kinase and cyclic 2,3-diphosphoglycerate synthetase from Methanothermus fervidus.” FEBS Lett 272(1-2): 94–98.

Lentzen, G. and T. Schwarz (2006a). Kompatible Solute: Mikrobielle Herstellung und Anwendung. Angwandte Mikrobiologie, Springer Berlin Heidelberg: Berlin, Heidelberg: 355–370.

Lentzen, G. and T. Schwarz (2006b). “Extremolytes: Natural compounds from extremophiles for versatile applications.” Applied microbiology and biotechnology 72: 623–634.

Lippert, K. and E. A. Galinski (1992). “Enzyme stabilization be ectoine-type compatible solutes: protection against heating, freezing and drying.” Applied Microbiology and Biotechnology 37(1): 61–65.

Liu, Y. F., N. Zhang, X. Liu, X. Wang, Z. X. Wang, Y. Chen, H. W. Yao, M. Ge and X. M. Pan (2012). “Molecular mechanism underlying the interaction of typical Sac10b family proteins with DNA.” PLoS One 7(4): e34986.

Martins, L. O. and H. Santos (1995). “Accumulation of Mannosylglycerate and Di-myo-Inositol-Phosphate by Pyrococcus furiosus in Response to Salinity and Temperature.” Appl Environ Microbiol 61(9): 3299–3303.

Matussek, K., P. Moritz, N. Brunner, C. Eckerskorn and R. Hensel (1998). “Cloning, sequencing, and expression of the gene encoding cyclic 2, 3-diphosphoglycerate synthetase, the key enzyme of cyclic 2, 3-diphosphoglycerate metabolism in Methanothermus fervidus.” J Bacteriol 180(22): 5997–6004.

McNicholas, S., E. Potterton, K. S. Wilson and M. E. Noble (2011). “Presenting your structures: the CCP4mg molecular-graphics software.” Acta Crystallogr D Biol Crystallogr 67(Pt 4): 386–394.

Reed, C. J., H. Lewis, E. Trejo, V. Winston and C. Evilia (2013). “Protein adaptations in archaeal extremophiles.” Archaea 2013: 373275.

Reeve, J. N. (1993). Structure and Organization of Genes. Methanogenesis: Ecology, Physiology, Biochemistry & Genetics J. G. Ferry, Springer US: Boston, MA: 493–526.

Rosano, G. and E. Ceccarelli (2009). “Rare codon content affects the solubility of recombinant proteins in a codon bias-adjusted Escherichia coli strain.” Microbial cell factories 8: 41.

Sauer, T. and E. A. Galinski (1998). “Bacterial milking: A novel bioprocess for production of compatible solutes.” Biotechnol Bioeng 57(3): 306–313.

Scholz, S., J. Sonnenbichler, W. Schäfer and R. Hensel (1992). “Di-myo-inositol-1,1’-phosphate: a new inositol phosphate isolated from Pyrococcus woesei.” FEBS Lett 306(2-3): 239–242.

Seely, R. J. and D. E. Fahrney (1983). “A novel diphospho-P,P’-diester from Methanobacterium thermoautotrophicum.” J Biol Chem 258(18): 10835–10838.

Shen, L., M. Kohlhaas, J. Enoki, R. Meier, B. Schönenberger, R. Wohlgemuth, R. Kourist, F. Niemeyer, D. van Niekerk, C. Bräsen, J. Niemeyer, J. Snoep and B. Siebers (2020). “A combined experimental and modelling approach for the Weimberg pathway optimisation.” Nature Communications 11(1): 1098.

Shima, S., D. A. Hérault, A. Berkessel and R. K. Thauer (1998). “Activation and thermostabilization effects of cyclic 2,3-diphosphoglycerate on enzymes from the hyperthermophilic Methanopyrus kandleri.” Arch Microbiol 170(6): 469–472.

van Alebeek, G.-J. W. M., G. Tafazzul, M. J. J. Kreuwels, J. T. Keltjens and G. D. Vogels (1994). “Cyclic 2,3-diphosphoglycerate metabolism in Methanobacterium thermoautotrophicum (strain ΔH): characterization of the synthetase reaction.” Archives of Microbiology 162(3): 193–198.

Vera, A., N. González-Montalbán, A. Arís and A. Villaverde (2007). “The conformational quality of insoluble recombinant proteins is enhanced at low growth temperatures.” Biotechnol Bioeng 96(6): 1101–1106.

Young, C. L., Z. T. Britton and A. S. Robinson (2012). “Recombinant protein expression and purification: a comprehensive review of affinity tags and microbial applications.” Biotechnol J 7(5): 620–634.

