## Supplementary Tables and Figures for "Establishment of an efficient one-step enzymatic synthesis of cyclic-2,3-diphosphoglycerate"

[5]Institute of Environmental Sciences, Hebrew University of Jerusalem, Rehovot 7610001, Israel
[#]Current affiliation

[6]Living Systems Institute, Faculty of Health and Life Sciences, University of Exeter, Stocker Road, Exeter EX4 4QD, UK.

[7]Corporate Innovation, Evonik Industries AG, Essen, Germany

[8]Department of Biochemistry, University of Stellenbosch, Matieland, South Africa; Molecular Cell Biology, Vrije Universiteit Amsterdam

*** Correspondence:**
Corresponding Authors




**Keywords: hyperthermophiles, archaea, stress response, compatible solutes, extremolytes, thermoprotection, 2-phosphoglycerate kinase, cyclic-2,3-diphosphoglycerate synthetase**

**Abbreviations: (cDPGS) cyclic-2,3-diphosphoglycerate synthetase – (2,3DPG) 2,3-diphosphoglycerate – (cDPG) cyclic 2,3-diphosphoglycerate – (2PG) 2-phosphoglycerate – (2PGK) 2-phosphoglycerate kinase**

### 1 Supplementary Figures and Tables

| **Primer** | **Sequence (5'-3')** |
|---|---|
| f-*Nco*I_*cdpgs* | tctaccatggggggcgagacaaaaa |
| r-*Bam*HI_ *cdpgs* | catcggatccttagcggttgttctt |
| f-p24-*cdpgs*-*Hind*III | gacgaagcttatgggtgaaactaaaaaaatg |
| r-p24- *cdpgs*-*Xho*I | gcacctcgagcctattgtttttaaaatcatctattgcc |
| f-*EcoR*I-*cdpgs* | tcgtgaattcatgaccgccgtgaagaggatac |
| r-*Hind*III-*cdpgs* | taataagcttaccctcgaaccgatcgatcgcca |
| **Plasmids/Strains** | **Source** |
| *pET15b* | Merck, Darmstadt, Germany |
| *pET24a* | Merck, Darmstadt, Germany |
| ***pEX_K4_cdpgs_Mfer_opt_bac*** | Eurofins genomics, Ebersberg, Germany |
| *pET24a_cdpgs_orig:His-tag* | This work |
| *pET15b_ cdpgs_opt:His-tag* | This work |
| *pET15b_cdpgs_opt_no-tag* | This work |
| *E. coli DH5α* | Life Technologies, Carlsbad, USA |
| ***E. coli BL21(DE3)-CodonPlus-pRIL (Cam$^R$)*** | Merck, Darmstadt, Germany |
| *E. coli BL21(DE3)* | Merck, Darmstadt, Germany |
| Rosetta(DE3) | Stratagene, San Diego, USA |

**Supplementary Table 1**: List of primers, plasmids and strains used in this study.

**Supplementary Table 2**: Sequence comparison of *cdpgs* gene from *M. fervidus* and the codon optimized version.

| ***Mfer* sequence (*Mfer*_0077, CAA70986)_*Mfcdpgs* original** |
|---|
| ATGGGTGAAACTAAAAAAATGATATGCTTAGTAGATGGTGAACATTATTTTCCTGTAGTTAAAGATT<br>CTATTGAAATATTGGATGATCTTGAACATATAGACGTTGTAGCTGTAGTATTCATTGGGGGAACTGA<br>AAAACTCCAAATTGAAGATCCTAAAGAATATTCTGAAAAATTAGGAAAACCAGTTTTTTTTGGGCCT<br>GATCCAAAAAAAATACCATATGATGTAATAAAAAAATGTGTAAAAAAATATAATGCTGATATAGTT<br>ATGGATTTAAGTGATGAACCTGTTGTAGATTATACAAAAAGATTTAGAATAGCATCCATAGTTCTAA<br>AAGAAGGCGCTGTATATCAGGGCGCTGATTTTAAATTTGAACCACTTACAGAATATGATGTGTTAGA<br>AAAGCCATCGATTAAAATTATAGGAACTGGAAAAAGAATAGGAAAAACTGCGGTATCTGCATATGC<br>TGCAAGAGTCATTCATAAACATAAATACAATCCATGTGTAGTTGCGATGGGTCGTGGAGGACCACGA<br>GAACCTGAAATAGTAGAAGGAAATAAAATAGAAATAACAGCTGAATATTTATTAGAACAAGCTGAT<br>AAAGGTGTACATGCAGCATCAGATCATTGGGAAGATGCATTAATGAGTAGAATTCTTACAGTCGGAT<br>GTAGAAGATGTGGCGGTGGAATGCTTGGTGATACATTCATAACAAATGTGAAAAGAGGTGCTGAAA<br>TTGCCAATAAATTAGATTCTGACTTTGTTATAATGGAAGGTAGTGGAGCAGCAATACCTCCTGTTAA<br>AACAAATAGGCAAATAGTTACAGTTGGTGCAAATCAGCCAATGATAAATATTAACAATTTCTTTGGA<br>CCATTTAGGATAGGATTAGCAGATCTTGTCATAATAACAATGTGTGAAGAACCTATGGCAACTACAG<br>AAAAAATTAAAAAGGTAGAAAAATTTATAAAAGAAATAAATCCATCAGCAAATGTTATTCCTACTGT<br>ATTTAGACCTAAACCAGTAGGCAATGTTGAAGGTAAAAAAGTGTTATTTGCAACTACAGCACCAAAA<br>GTTGTTGTAGGGAAATTAGTTAATTATCTAGAAAGTAAATATGGATGCGATGTGGTAGGTGTCACAC<br>CACATTTATCAAATCGTCCATTATTACGTAGAGATTTAAAAAAATATATAAACAAGGCAGATTTAAT<br>GTTGACGGAATTAAAGGCTGCAGCTGTTGACGTTGCAACTAGGGTAGCTATAGAAGCTGGTCTAGAT<br>GTTGTATATTGTGATAATATTCCTGTAGTTATAGATGAGAGTTATGGAAACATCGATGATGCAATTAT<br>TGAAGTTGTAGAAATGGCAATAGATGATTTTAAAAACAATAGGTGA |
| ***Mfcdpgs* codon optimized for expression in *E. coli* (Eurofins Genomics, Ebersberg, Germany)** |
| ATGGGCGAGACAAAAAAGATGATTTGCCTGGTAGATGGGGAACACTATTTTCCTGTTGTTAAAGACA<br>GCATTGAAATCCTCGATGATCTGGAGCATATCGACGTAGTGGCTGTGGTATTCATCGGCGGAACCGA<br>GAAACTGCAGATTGAAGATCCGAAAGAATATTCGGAAAAACTGGGCAAACCTGTGTTCTTTGGACCC<br>GATCCGAAGAAAATTCCGTATGACGTTATCAAGAAATGCGTCAAGAAATACAATGCGGATATTGTGA<br>TGGATCTTTCTGACGAACCAGTAGTGGACTACACCAAACGGTTTCGCATCGCCTCCATTGTGCTGAA<br>AGAGGGCGCAGTTTATCAAGGGGCCGATTTTAAATTTGAACCGCTGACTGAATACGATGTTTTGGAG<br>AAACCGTCTATCAAAATTATTGGTACCGGGAAACGCATTGGTAAGACAGCGGTGAGTGCGTATGCAG<br>CCCGTGTGATTCACAAGCATAAATACAATCCCTGTGTAGTTGCAATGGGCCGTGGTGGACCACGTGA<br>ACCGGAGATTGTGGAGGGCAACAAAATCGAAATCACCGCCGAATATCTGCTTGAGCAAGCGGATAA<br>AGGCGTTCATGCAGCCAGCGATCATTGGGAAGATGCCCTGATGAGTCGCATTCTGACGGTTGGATGT<br>CGTCGTTGTGGTGGTGGGATGCTGGGCGACACGTTCATTACCAACGTCAAACGTGGTGCAGAGATTG<br>CGAACAAACTGGACTCAGATTTTGTCATTATGGAAGGTTCAGGTGCGGCAATTCCGCCGGTGAAAAC<br>GAATCGGCAGATTGTCACTGTTGGCGCCAATCAGCCGATGATCAACATCAATAACTTCTTTGGCCCG<br>TTTCGCATTGGCTTAGCCGATTTGGTCATCATTACCATGTGTGAAGAACCGATGGCGACCACCGAAA<br>AGATCAAGAAAGTTGAGAAATTCATTAAAGAGATCAATCCCAGCGCTAATGTGATTCCGACGGTTTT<br>CCGCCCAAAACCTGTGGGTAACGTCGAAGGTAAAAAAGTGTTGTTTGCGACCACGGCCCCGAAAGTT<br>GTGGTAGGGAAACTCGTGAATTACCTGGAATCGAAATATGGCTGCGATGTAGTGGGTGTTACGCCAC<br>ACCTGAGCAATCGCCCTCTGTTACGTCGCGATTTAAAGAAATACATTAACAAAGCGGATCTTATGCT<br>CACTGAACTGAAAGCGGCTGCTGTGGATGTCGCGACACGCGTAGCTATTGAAGCGGGCTTAGATGTC<br>GTGTATTGCGACAACATCCCAGTCGTCATCGACGAATCCTATGGCAACATTGACGATGCAATCATCG<br>AAGTGGTCGAAATGGCTATCGACGACTTCAAGAACAACCGCTAA |
| **Comparison of native- versus codon optimized-sequence (differences highlighted in green)** |

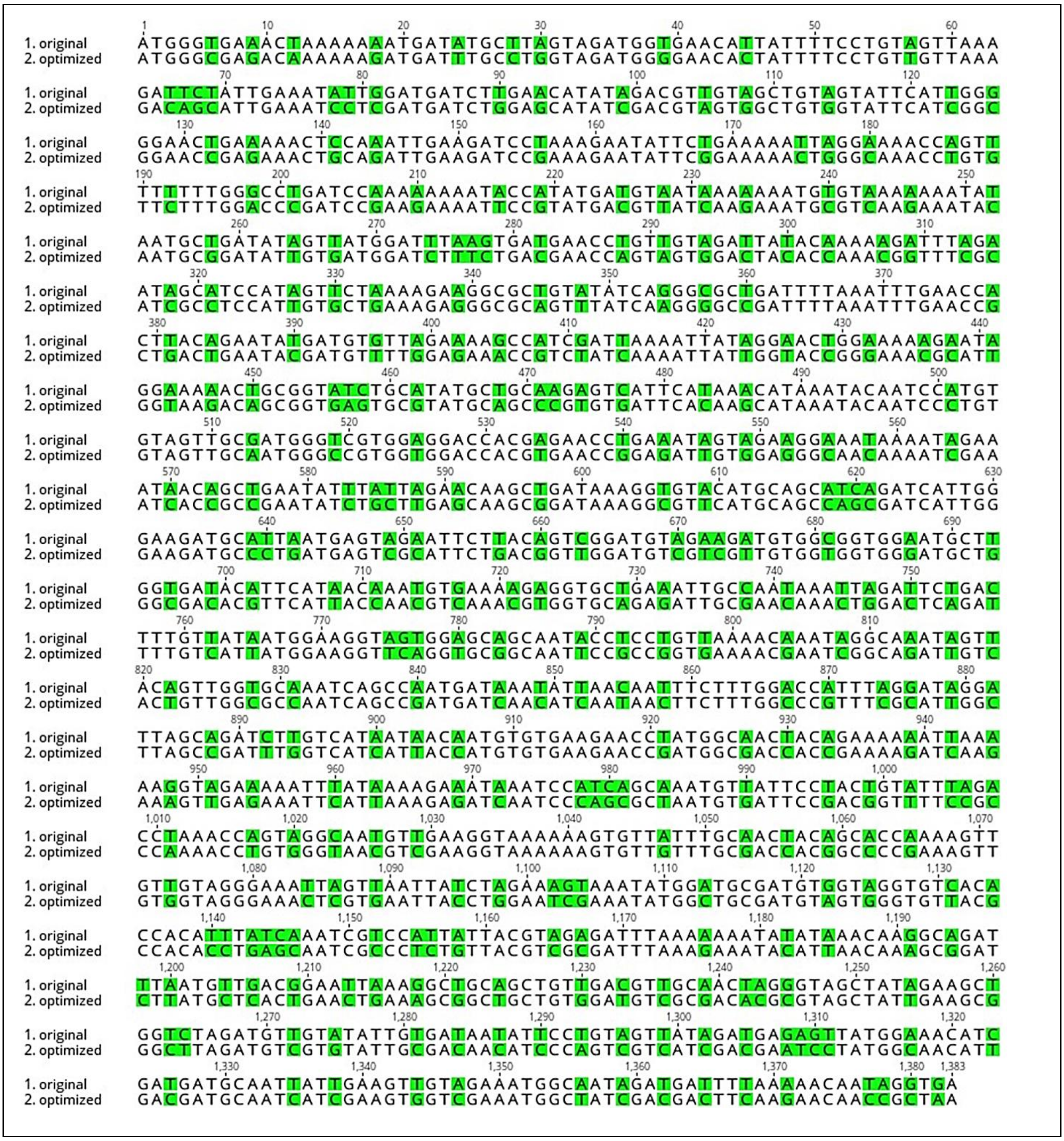
1. original
2. optimized

**Supplementary Table 3**: Comparison of purification state, assay conditions and kinetic parameters of *Mf*cDPGS.

| cDPG synthesis | *M. fervidus* | | *E. coli* | | *E. coli* | |
|---|---|---|---|---|---|---|
| | *(Matussek et al., 1998)* | | *(Matussek et al., 1998)* | | *(this study)* | |
| Purification grade: | Reactive green agarose | | Reactive green agarose | | Size exclusion chromatography | |
| Assay conditions: | 10 mM TES/$K^+$ (pH 7.0), 875 mM KCl, 5 mM DTT, 83°C, $MgCl_2$ = *NA | | 10 mM TES/$K^+$ (pH 7.0), 875 mM KCl, 5 mM DTT, 83°C, $MgCl_2$ = *NA | | 50 mM MES/KOH (pH 6.5), 400 mM KCl, 10 mM DTT, 55°C, $MgCl_2$ = 10 mM | |
| Substrate: | $K_m$ | $V_{max}$ | $K_m$ | $V_{max}$ | $K_m$ | $V_{max}$ |
| **2,3 DPG** | 5.8 ±2.0 | 32.8 ±4.0 | 5.9 ±2.5 | 17.2 ±2.0 | 1.52 ±0.40 | 38.2 ±1.70 |
| **ATP** | 3.0 ±1.5 | 32.0 ±3.0 | 3.0 ±1.5 | 16.0 ±2.0 | 0.55 ±0.08 | 38.2 ±1.70 |

NA= not annotated in the work of Matussek and coworkers

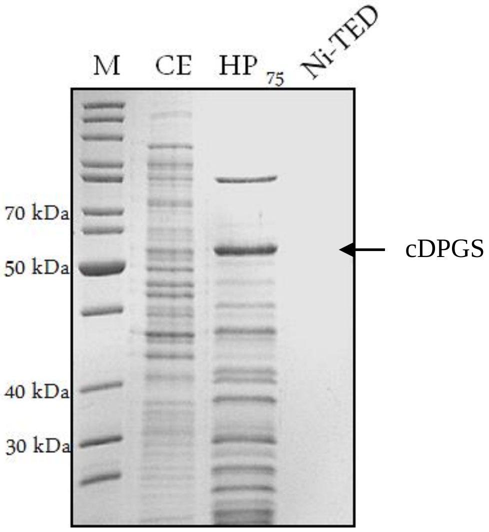


**Supplementary Figure S1: Heterologous expression and purification of recombinant *Mf*cDPGS** C-terminal His-tag in *E. coli*. The *cdpgs* gene was cloned in pET24a with C-terminal His-tag and expressed in *E. coli* BL21(DE3)-Codon-Plus. The protein fractions after successive purification steps were analyzed via SDS-PAGE (12.5 %) and stained with Coomassie Brilliant Blue. M: Protein marker, unstained protein ladder (Thermo Fisher Scientific, Carlsbad, USA), CE: Crude extract (5 µg), HP: Soluble fraction after heat precipitation (75°C, 20 min, 5 µg), Ni-TED: elution fraction after Ni-TED affinity chromatography. The recombinant cDPGS with C-terminal His-tag does not bind to the column. The arrow indicates the expected size of cDPGS (50.7 kDa).

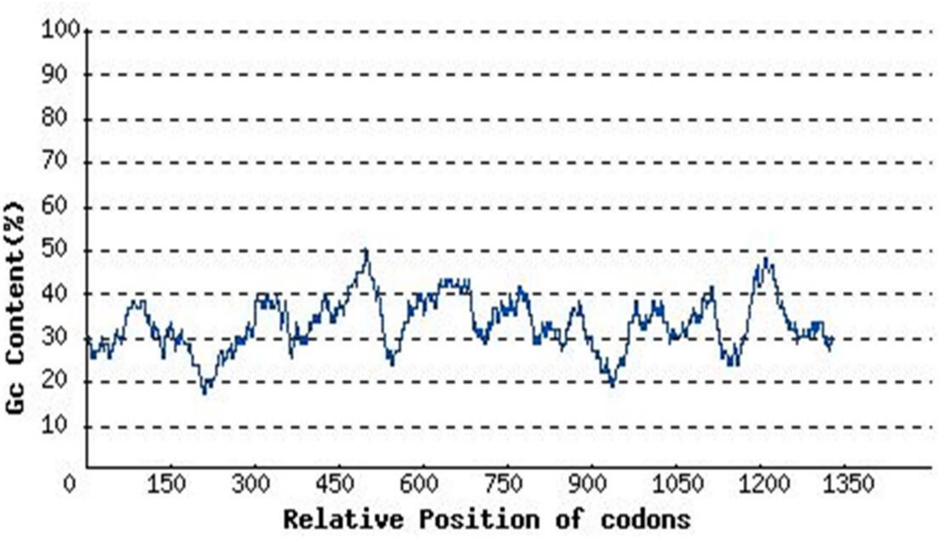


**Supplementary Figure S2: GC-content analysis of *Mf*cDPGS.** The average GC-content was calculated to be 32.8 % and the distribution of the of GC-content is mapped over the whole gene sequence using the genescript rare codon analysis tool (https://www.genscript.com/tools/rare-codon-analysis).

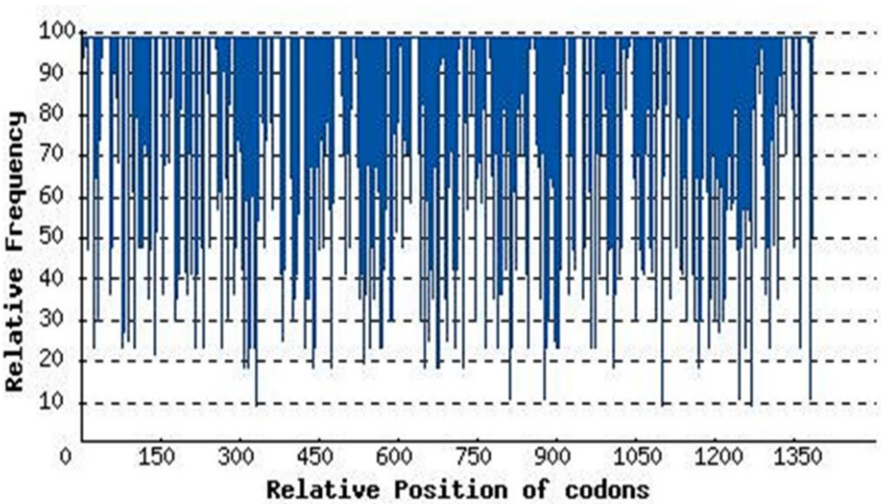


**Supplementary Figure S3: Codon-adaption-index of the *Mf*cDPGS.** The relative codon frequency for the desired expression host *E. coli* is mapped over the whole gene sequence of the *cdpgs*. The final calculated resulted was a codon adaption index of the *cdpgs* for the expression in *E. coli* of 0.54. The analysis was performed using the genescript rare codon analysis tool (https://www.genscript.com/tools/rare-codon-analysis).

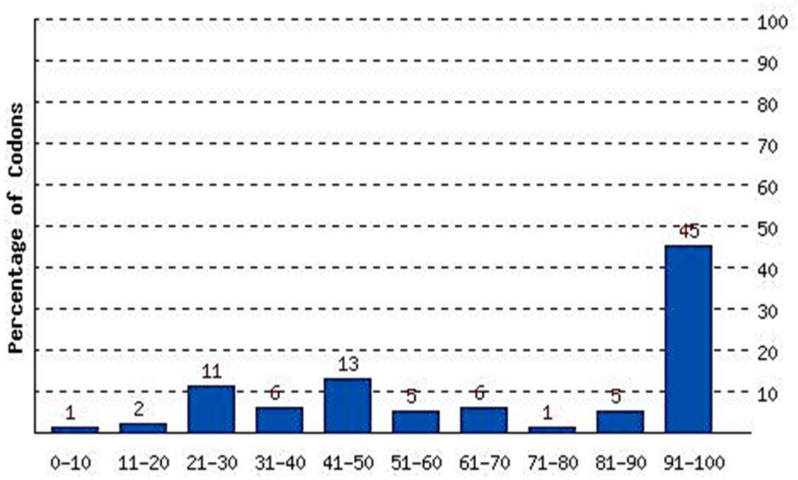


**Supplementary Figure S4: Codon frequency distribution of the *Mf*cDPGS..** The percentage distribution was calculated in computed quality groups and compared to that of the desired expression host *E. coli*. The result showed a total number of 14 codons with lower values than 31%, which will lower the the expression efficiency significantly. The analysis was performed using the genescript rare codon analysis tool (https://www.genscript.com/tools/rare-codon-analysis).

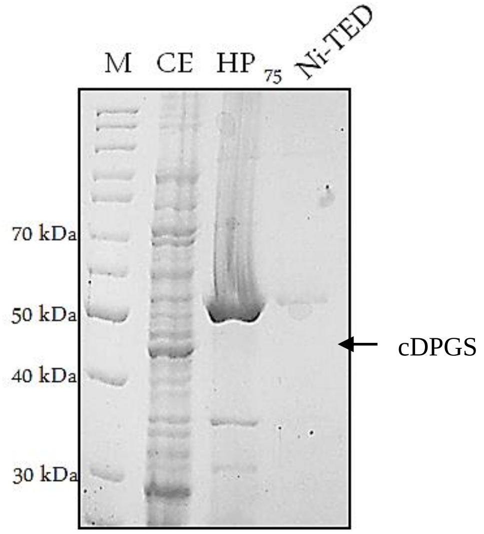


**Supplementary Figure S5: Expression and purification of recombinant codon optimized *Mf*cDPGS with N-terminal His-tag.** The codon optimized *cdpgs* gene was cloned in pET15b with N-terminal His-tag and expressed in *E. coli* BL21(DE3)-Codon-Plus. The protein fractions after successive purification steps were analyzed via SDS-PAGE (12.5 %) and stained with Coomassie Brilliant Blue. M: Protein marker, unstained protein ladder (Thermo Fisher Scientific, Carlsbad, USA), CE: Crude extract (5 µg), HP: Soluble fraction after heat precipitation (75°C, 30 min, Eppendorf Thermomixer at 700 rpm, 5 µg), Ni-TED: elution fraction after Ni-TED affinity chromatography (small quantities of N-terminal could be purified, 0.8 µg). The arrow indicates the expected size of cDPGS (50.7 kDa).

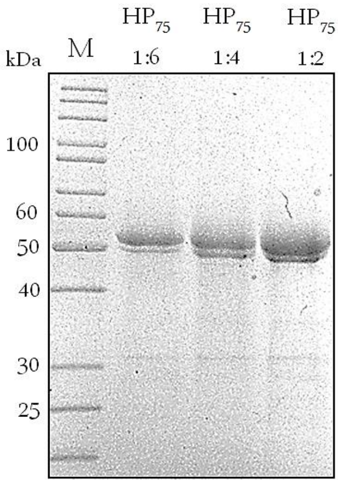


**Supplementary Figure S6: Optimization of heat purification of the recombinant codon optimized *Mf*cDPGS without affinity tag from *M. fervidus*.** The codon optimized *cdpgs* was cloned in pET15b without affinity tag and expressed in *E. coli* BL21(DE3)-Codon-Plus. The obtained crude extract was diluted (1:2, 1:4, 1:6) in MES/KOH, pH 6.5, 400 mM KCl, 10 mM $MgCl_2$, 10 mM DTT and subjected to a heat precipitation step at 75 °C for 30 min in an Eppendorf Thermomixer at 700 rpm. Protein fractions after heat purification steps were analyzed via SDS-PAGE (12.5 %) and stained with Coomassie Brilliant Blue. M: Protein marker, unstained protein ladder (Thermo Fisher Scientific, Carlsbad, USA). HP: heat precipitation fractions (5 µg) are shown.

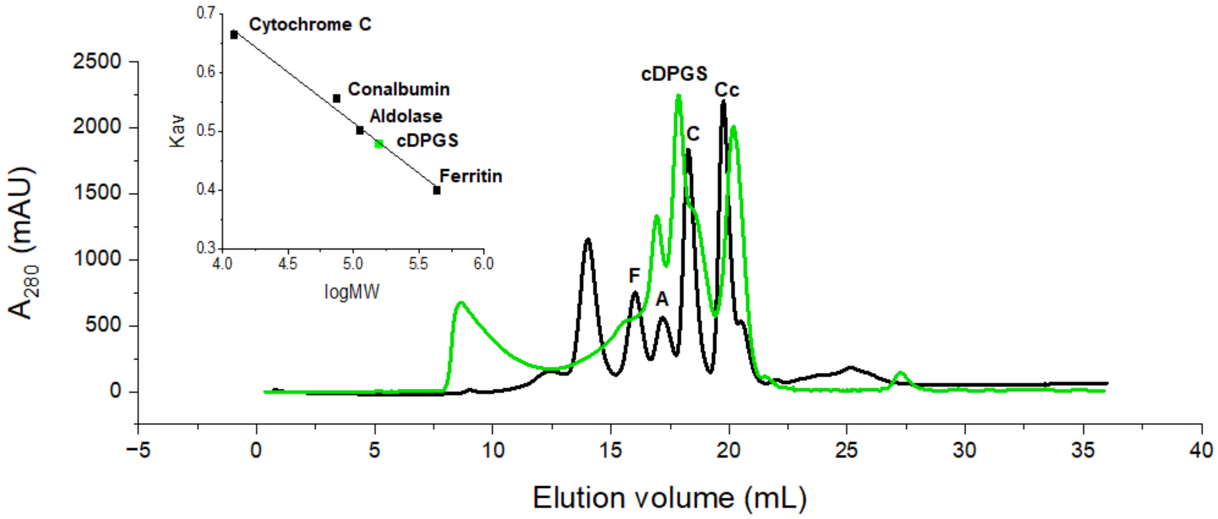


**Supplementary Figure S7: Size exclusion chromatography of the recombinant *Mf*cDPGS.** The *Mf*cDPGS after heat precipitation at 75°C (5 mg) was applied onto a Superose Increase 10/300 column, equilibrated in 50 mM MES/KOH, 400 mM KCl, pH 6.5, flow rate 0.5 mL/min. In the elution profiles of the *Mf*cDPGS is shown in green and the standard proteins for calibration curve are shown in black. Abbreviations represent the following marker proteins: (F) – Ferritin, (A) – Aldolase, (C) – Conalbumin, (Cc) - Cytochrom C. The standard K average – log MW graph for the estimation of the molecular weight of the *Mf*cDPGS is shown as inlay in the upper left corner. The native molecular mass was 119 kDa confirming the homodimeric structure of the protein.

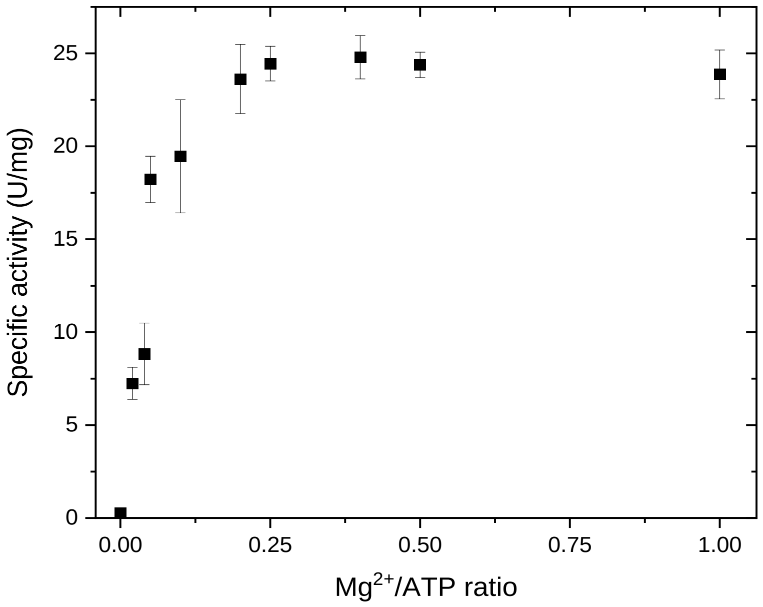


**Supplementary Figure S8: Activity of recombinant *Mf*cDPGS as function of the $Mg^{2+}$/ATP ratio.** The activity was measured using the PK-LDH assay. The assays were performed in presence of different molar ratios of $Mg^{2+}$/ATP ranging from 0.02 – 1.0 (1 – 50 mM) under optimized assay conditions (50 mM MES/KOH pH 6.5, 400 mM KCl, 10 mM DTT, 10 mM 2,3 DPG) using 4.8 µg purified *Mf*cDPGS. The experiments were performed in technical triplicates, the error bars indicate the standard deviation of the mean.

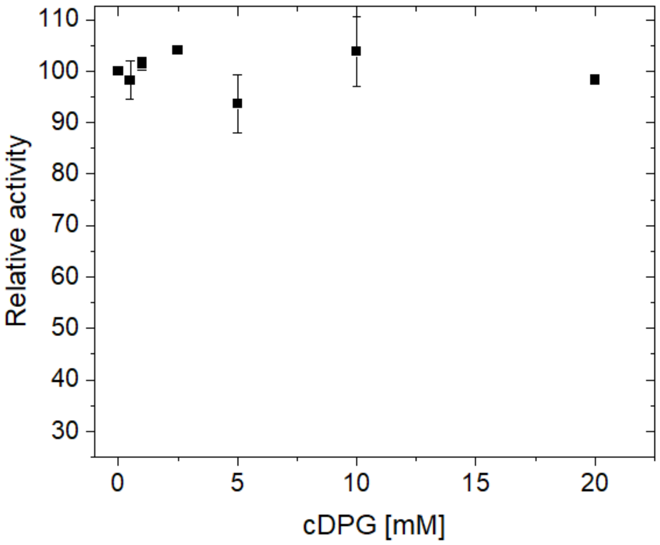


**Supplementary Figure S9: Product inhibition of cDPG on activity of the recombinant *Mf*cDPGS.** The activity was measured using the PK-LDH assay. The assays were performed in presence of various cDPG concentrations (0-20 mM) under optimized assay conditions (50 mM MES/KOH pH 6.5, 400 mM KCl, 10 mM DTT, 5 mM 2,3DPG, 10 mM ATP, 20 mM $MgCl_2$ ($Mg^{2+}$/ATP 0.5), 55°C) using 4.8 µg purified cDPGS. The experiments were performed in technical duplicates (since only limited amounts of cDPG were available), the error bars indicate the standard deviation of the mean. 100 % of relative activity corresponds to a specific activity of 24.23 U $mg^{-1}$.

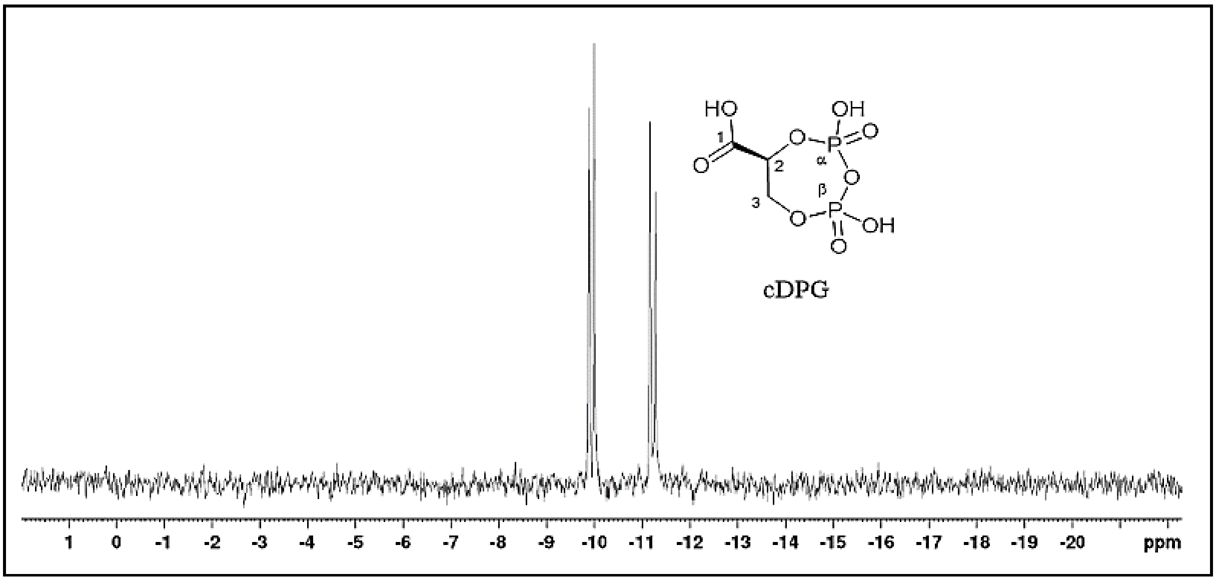


**Supplementary Figure S10: $^{31}$P-NMR analysis of cDPG.** 5 mM cDPG was solved in 50 mM MES/KOH, pH 6.5 supplemented with 400 mM KCl, 10 mM $MgCl_2$, and 20% (v/v) $D_2O$. The $^{31}$P-NMR spectra was acquired using a Bruker Avance Neo 400 spectrometer (400MHz, 80 % buffered sample and 20 % $D_2O$) δ[ppm] = -9.94 (d, $^{3}J$□,□=18.1 Hz, 1P, □-P), -11.21 (d, $^{3}J$□,□=18.1 Hz, 1P, □-P). Data processing was performed using TopSpin© 3.6.2 software (Bruker BioSpin, Rheinstetten, Germany).

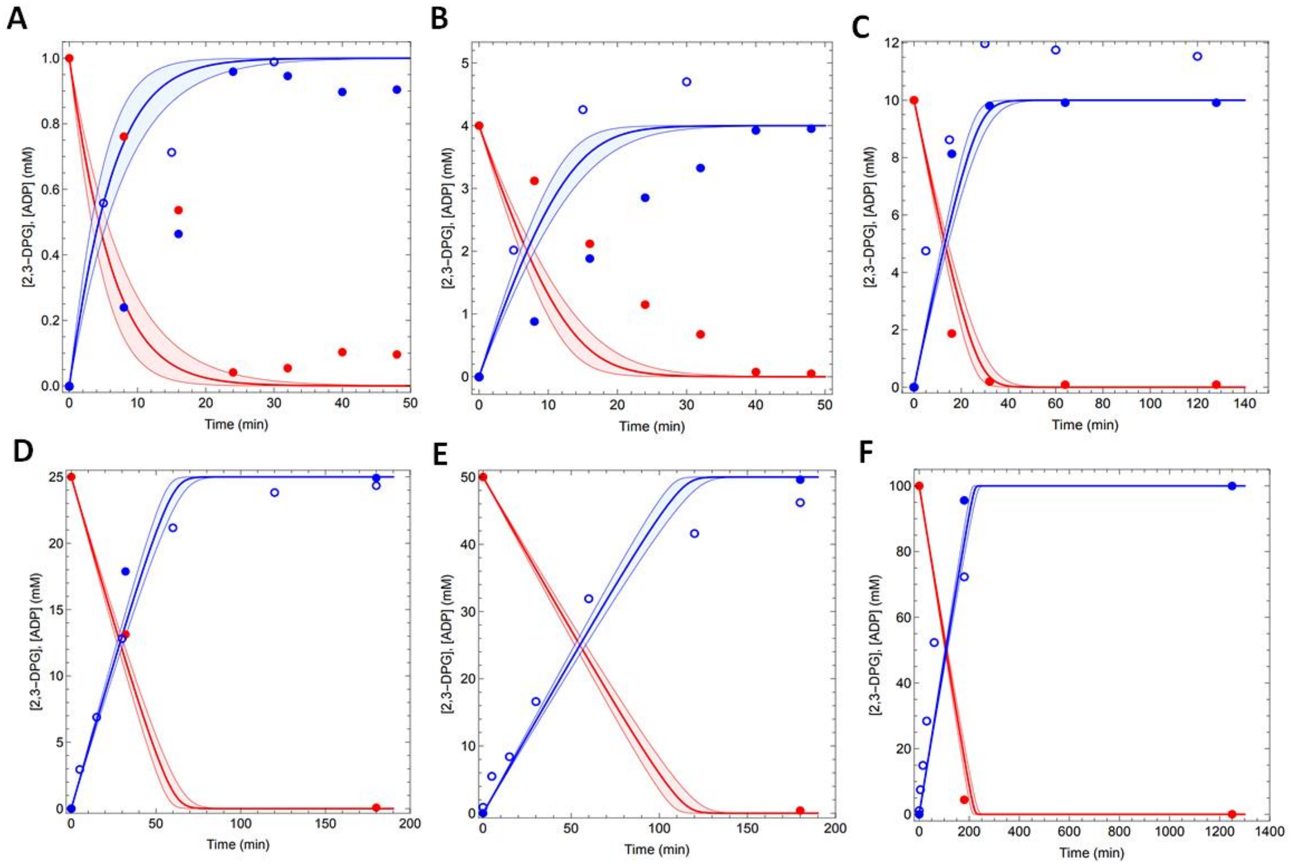


**Supplementary Figure S11: Experimental data combined with mathematical model simulation for the time dependent conversion of 2,3 DPG to cDPG via recombinant *Mf*cDPGS.** The time-dependent conversion results for (**A**) 1 mM, (**B**) 4 mM, (**C**) 10 mM, (**D**) 25 mM, (**E**) 50 mM, and (**F**) 100 mM 2,3 DPG to cDPG are shown. The blue open circles represent the experimental data for the observed ADP concentrations, measured discontinuously. Assays were performed at 55°C in 50 mM MES/KOH (pH 6.5), 400 mM KCl and a $Mg^{2+}$/ATP ratio of 0.2 (e.g., 10 mM $MgCl_2$ / 50 mM ATP) using 0.5 U of purified *Mf*cDPGS. Samples were taken at regular time intervals, and the formed ADP was quantified using the continuous PK-LDH assay. The red filled circles represent the 2,3 DPG consumption, while the blue filled circles represent the cDPG concentrations, both quantified via $^{31}$P-NMR spectroscopy. Each data point represents the mean value of three independent technical replicates. The model predictions (eq. 1 and 2 main manuscript) for the experiments are shown with a red line for 2,3-DPG consumption and a blue line for cDPG production.

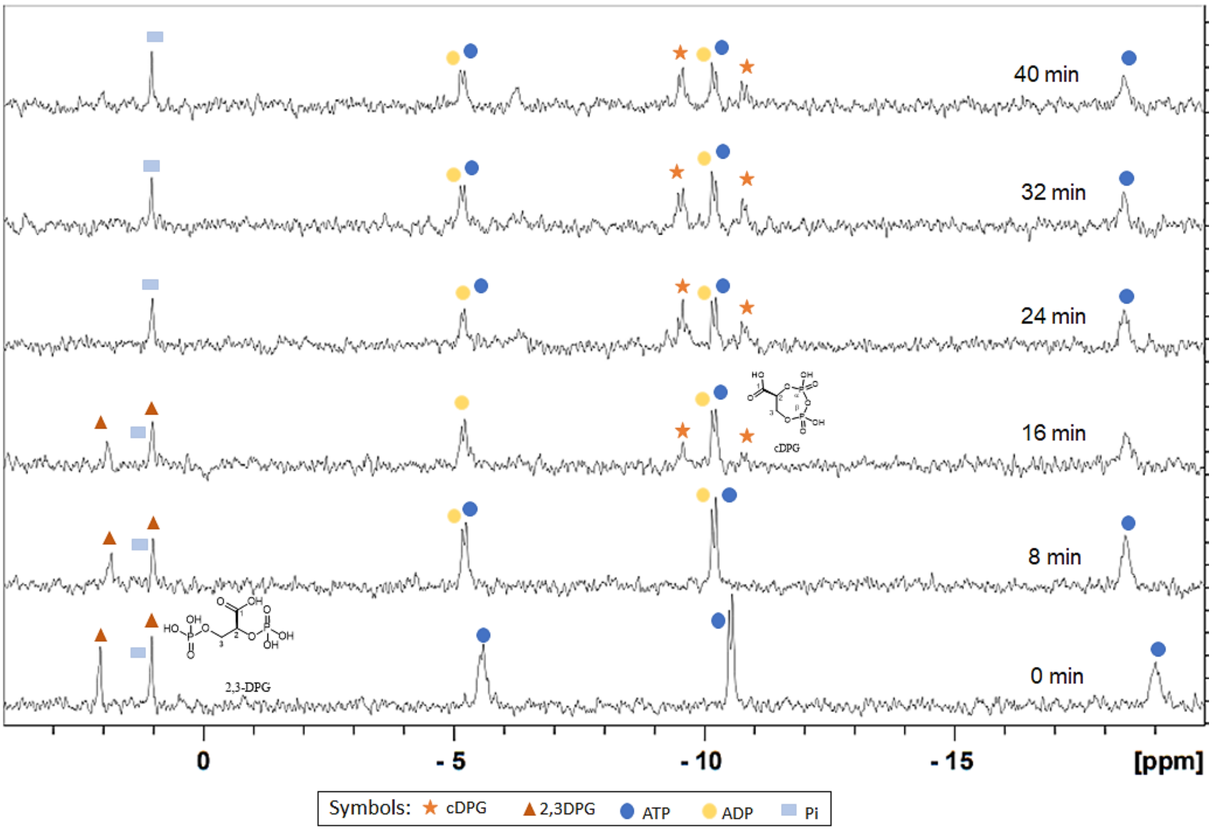
40 min
32 min
24 min
16 min
8 min
0 min
cDPG
2,3-DPG
0
- 5
- 10
- 15
[ppm]
Symbols: cDPG 2,3DPG ATP ADP Pi

6 A

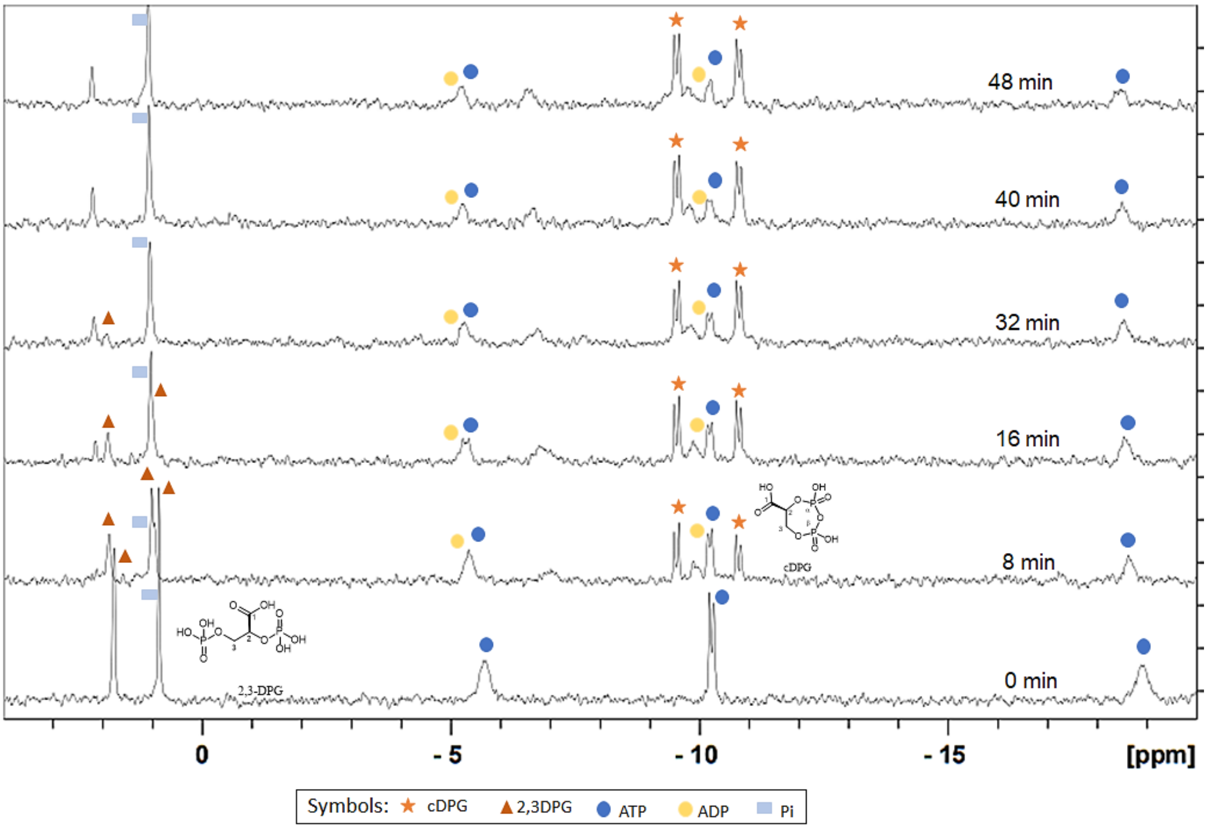
48 min
40 min
32 min
16 min
8 min
0 min
cDPG
2,3-DPG
0
- 5
- 10
- 15
[ppm]
Symbols: cDPG 2,3DPG ATP ADP Pi

6 B

6 C

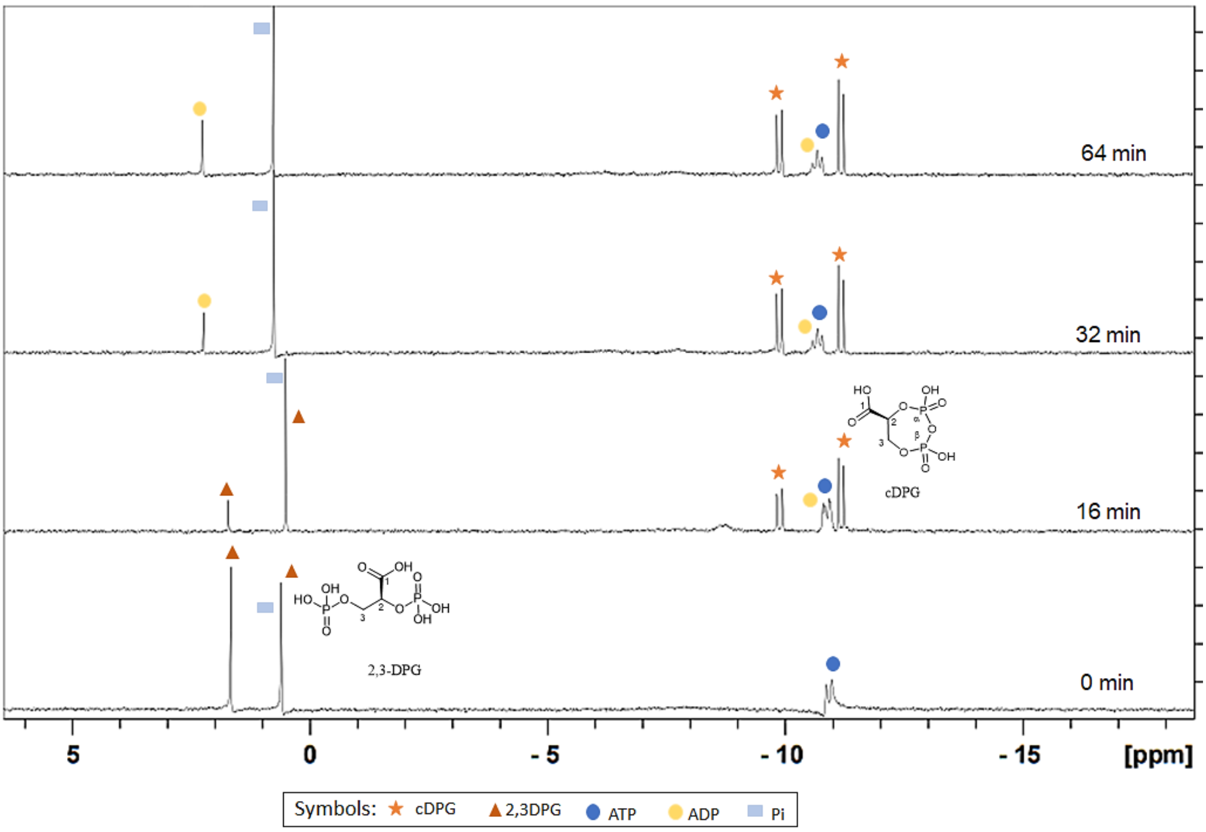
64 min
32 min
16 min
0 min
cDPG
2,3-DPG
5
0
- 5
- 10
- 15
[ppm]
Symbols: cDPG 2,3DPG ATP ADP Pi

6 D

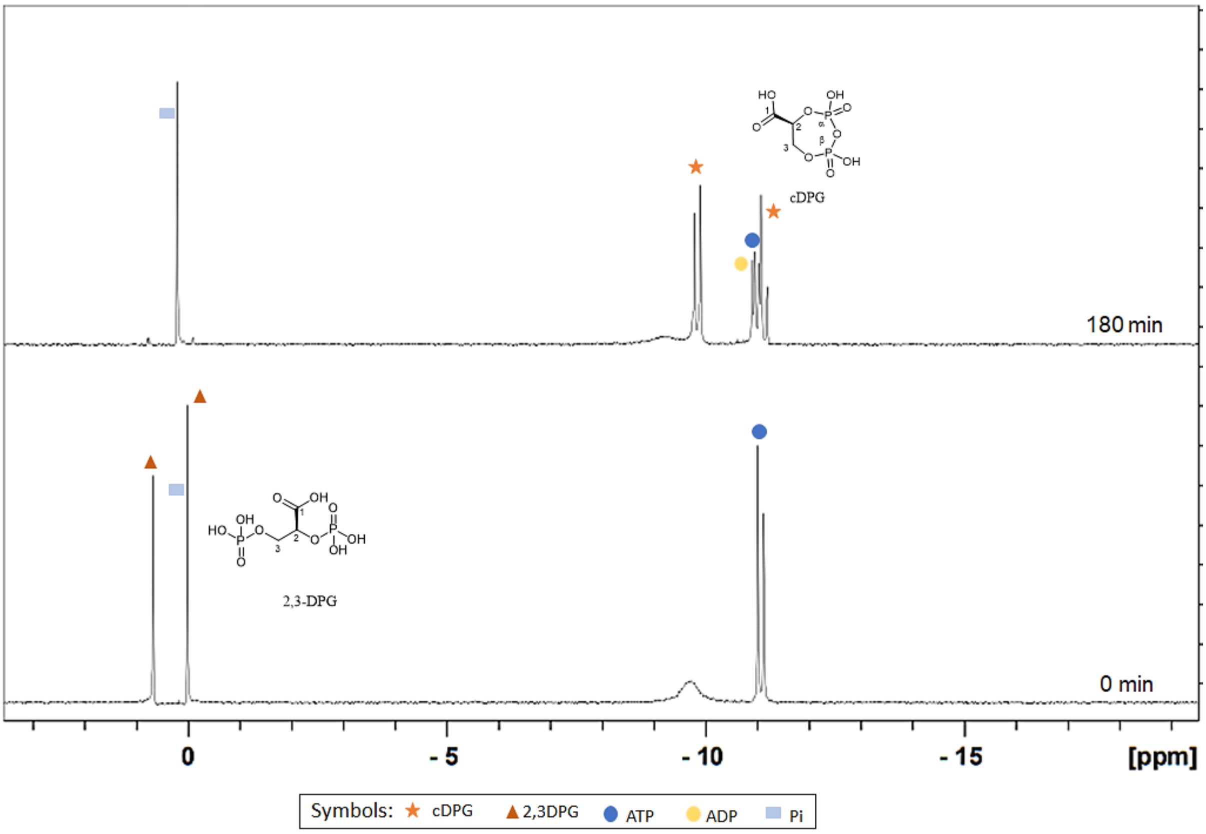
180 min
0 min
cDPG
2,3-DPG
0
- 5
- 10
- 15
[ppm]
Symbols: cDPG 2,3DPG ATP ADP Pi

6 E

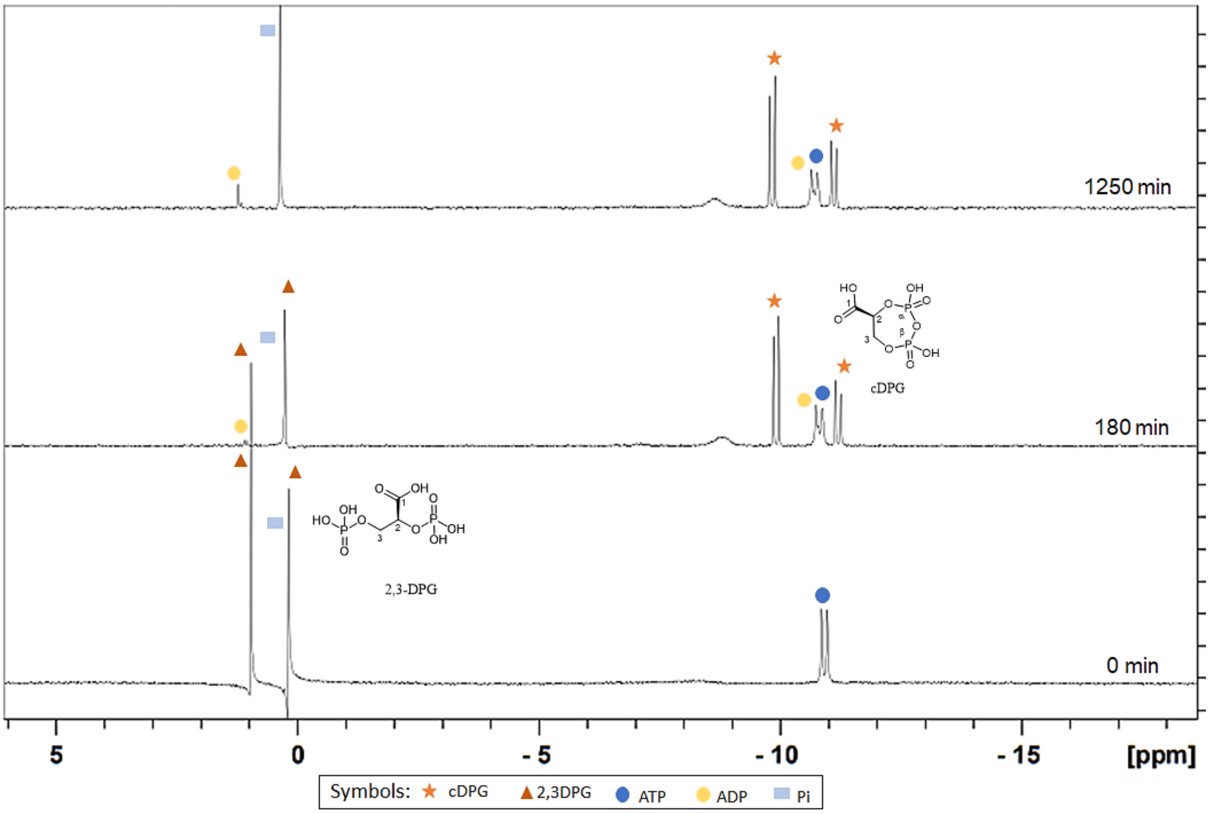


6 F

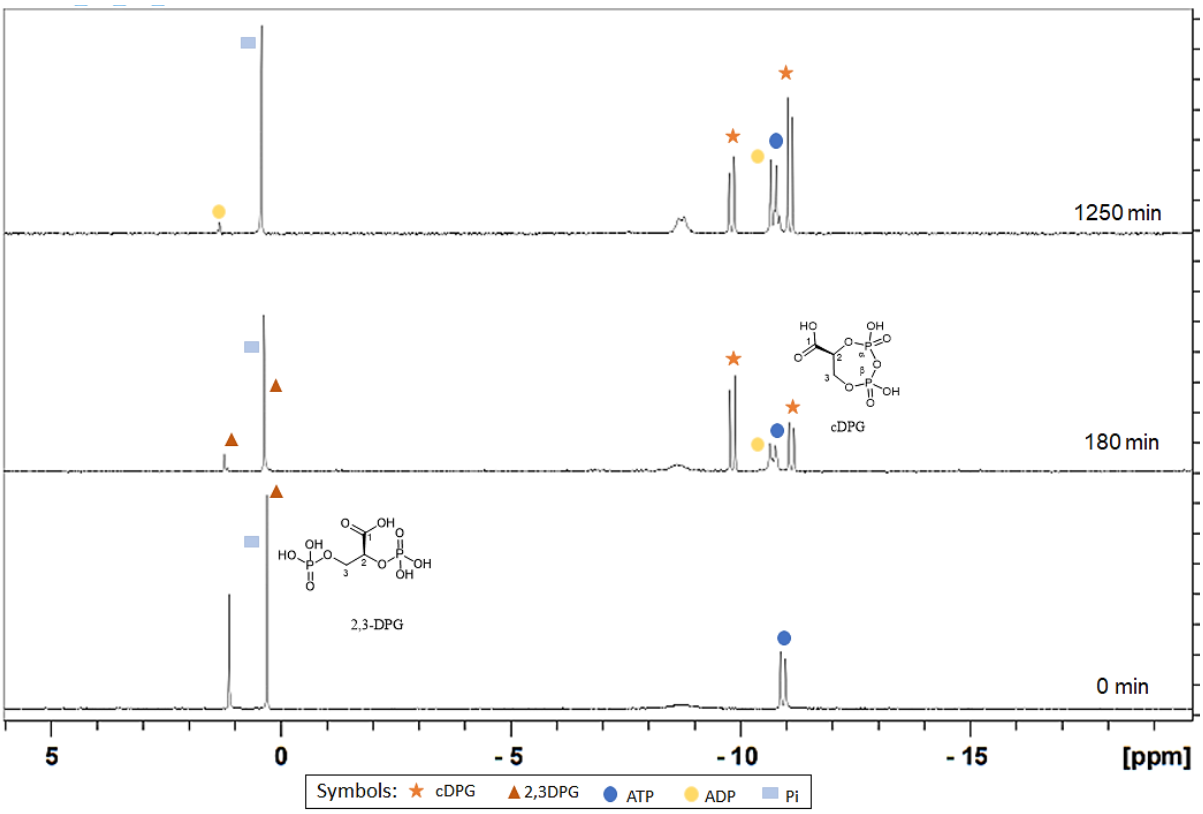


**Supplemental Figure S12 (A-F): Time-dependent conversion of 2,3 DPG to cDPG by *Mf*cDPGS, analyzed via $^{31}$P-NMR spectroscopy.** Conversion assays were performed using a concentration range of 1 – 100 mM 2,3 DPG. The corresponding stacked $^{31}$P-NMR spectra are shown for the time depended conversion of (**A**) 1 mM, (**B**) 4 mM, (**C**) 10 mM, (**D**) 25 mM, (**E**) 50 mM, and (**F**) 100 mM 2,3 DPG. All assay was performed discontinuously in 1 mL total volume at 55°C in 50 mM MES/KOH (pH 6.5), 400 mM KCl, and a $Mg^{2+}$/ATP ratio of 0.2 (e.g., 10 mM

$MgCl_2$ / 50 mM ATP) using 0.5 U of purified *Mf*cDPGS after SEC. Samples from the corresponding time intervals were mixed with $D_2O$ (20% v/v) and $^{31}P$ spectra were acquired using a Bruker Avance Neo 400 spectrometer (Bruker, Rheinstetten, Germany). Data processing was conducted using TopSpin© 3.6.2 software (Bruker BioSpin, Rheinstetten, Germany). The following species were identified in the spectra: cDPG (orange asterisk), 2,3-DPG (dark orange triangle), ATP (blue circle), ADP (yellow circle), and inorganic phosphate ($P_i$) (light blue rectangle).

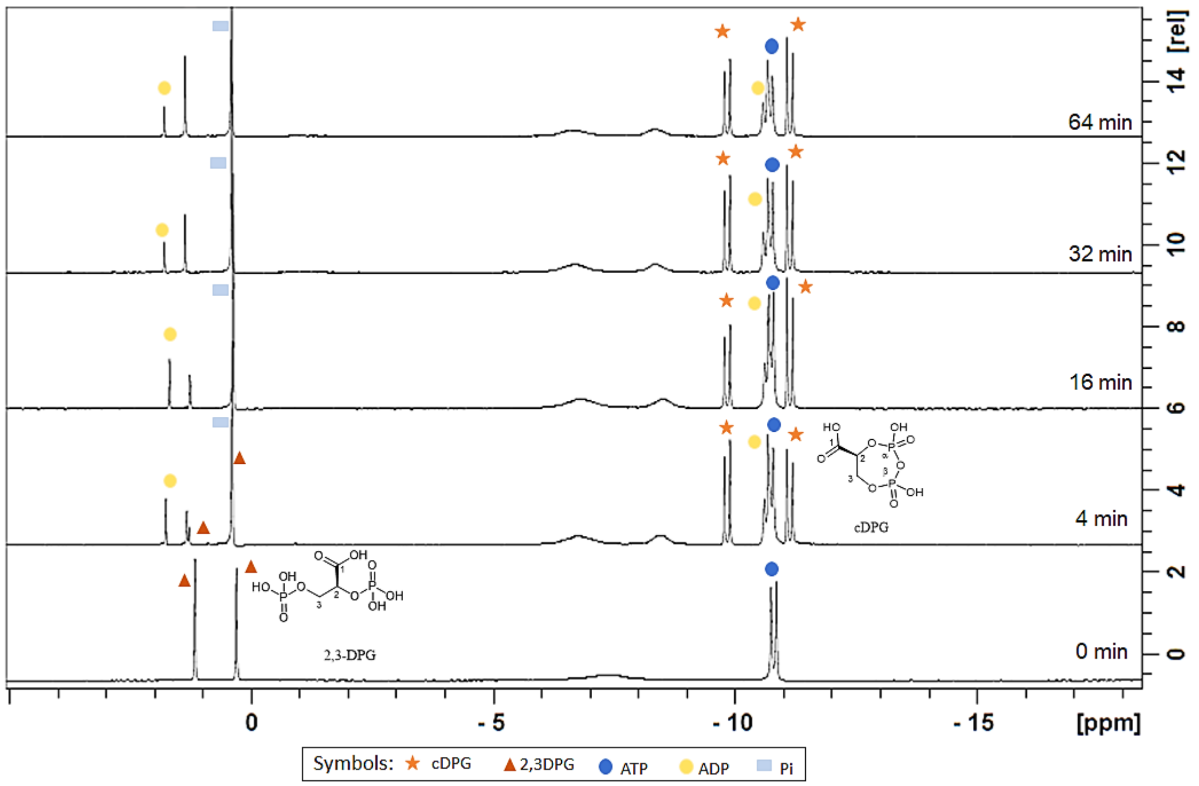


**Supplemental Figure S13: Time-dependent conversion of 10 mM 2,3 DPG to cDPG by heat purified *Mf*cDPGS analyzed via $^{31}$P-NMR.** A discontinuous enzyme assay using the heat purified cDPGS (0.65 U/ml) was performed at 55°C under optimized assay conditions (50 mM MES/KOH, pH 6.5 supplemented with 400 mM KCl, 10 mM DTT, 10 mM 2,3DPG, 20 mM ATP, $Mg^{2+}$/ATP 0.5
(10 mM $MgCl_2$/ 20 mM ATP) at 55°C, using 0.5 U purified *Mf*cDPGS). Samples were withdrawn at regular time intervals and mixed with $D_2O$ (20% v/v). $^{31}$P-NMR spectra were acquired using a Bruker Avance Neo 400 spectrometer. The stacked spectra view for respective time points are presented. Data processing was performed using TopSpin© 3.6.2 software (Bruker BioSpin, Rheinstetten, Germany). In the spectra, the following species were identified: orange asterisk represents cDPG, dark orange triangle represents 2,3 DPG, blue circle represents ATP, yellow circle represents ADP, and light blue rectangle represents inorganic phosphate ($P_i$).
